# Emergence of function from single RNA sequences by Darwinian evolution

**DOI:** 10.1101/2021.03.03.433769

**Authors:** Falk Wachowius, Benjamin T. Porebski, Christopher M. Johnson, Philipp Holliger

## Abstract

The spontaneous emergence of function from pools of random sequence RNA is widely considered an important transition in the origin of life. However, the plausibility of this hypothetical process and the number of productive evolutionary trajectories in sequence space are unknown. Here we demonstrate that function can arise starting from a single RNA sequence by an iterative process of mutation and selection. Specifically, we describe the discovery of both specific ATP or GTP aptamers - with micromolar affinity for their nucleotide ligand - starting each from a single, homopolymeric poly-A sequence flanked by conserved primer binding sites. Our results indicate that the *ab initio* presence of large, diverse random sequence pools is not a prerequisite for the emergence of functional RNAs and that the process of Darwinian evolution has the capacity to generate function even from single, largely unstructured RNA sequences with minimal molecular and informational complexity.

## Introduction

Multiple lines of evidence from modern biology support the hypothesis of an “RNA world”, a primordial biology where RNA played a more central role. These include, RNA nucleotide cofactors in present day metabolism, the central role of RNA in genetic processes, RNA catalysis in splicing, RNA processing and translation - i.e. the ribozyme cores of the spliceosome, RNAse P and ribosomal peptidyl-transferase centre^1^. Together with the discovery of varied and robust prebiotic synthesis routes to RNA (and DNA) nucleosides^2, 3^, their activation and efficient non-enzymatic polymerization suggest an increasingly plausible path towards the emergence of an RNA-based biology^4^.

A key transition along this path is thought to have been the spontaneous non-enzymatic polymerization of activated RNA nucleotides into pools of random sequence RNA oligomers from which functional sequences could emerge. The generation of functional sequences from random pools has been recapitulated and studied by *in vitro* selection experiments^5^, which have succeeded in the isolation of a diverse array of ligands (aptamers) and catalysts directly from random sequence pools of RNA, DNA and some unnatural nucleic analogues (XNA)^6^. Of particular interest in this context are nucleotide binding aptamers, which were among the first RNA aptamers to be isolated and which have been studied extensively as model systems. Such studies have sought to define the necessary and optimal size for binding as well as the sequence parameters necessary for aptamer discovery such as oligomer length, conformational pre-organization, pool complexity and adaptive landscapes^7–9^. Extrapolation from such selection experiments and analysis of the mutational landscapes of functional oligomers suggests that the frequency of distinct ligand-binding motifs is low (<10^-11^)^5^, even in complex random sequence pools, although structural pre-organization can increase the chance of ligand discovery^9^.

However, all of these experiments initiate from best-case scenarios regarding the size and sequence complexity of prebiotic oligomer pools, utilizing large, random sequence pools composed of approximately equal proportions of all four RNA nucleotides. However, studies on *ab initio* non-enzymatic polymerization of activated RNA nucleotides indicate very significant biases in plausible prebiotic pools^10–13^ with negative impacts on sequence complexity and presumably phenotypic potential, i.e. the probability of functional sequences in a given sequence pool. Indeed, fundamental studies on ligase ribozymes suggest that function (i.e. catalysis) may strongly scale with sequence complexity^14^. These findings thus pose a challenge to the conjecture of spontaneous emergence. The probability of the latter would seem to depend critically on pool size and sequence complexity, as these parameters inform both the initial frequency of functional sequences and the ease (the number of evolutionary trajectories) by which functional sequences can be accessed from arbitrary points in sequence space.

Here we critically test these arguments by exploring an extreme scenario of minimal pool complexity: the emergence of functional RNAs from a single, non-functional, RNA sequence. We performed five independent selection experiments starting each from a single, but distinct homopolymeric RNA sequence (comprising a central 39 nt poly-A or poly-U stretch flanked by different (in each case) and conserved 15 nt PCR primer binding sites) comprising cycles of error prone replication and selection for ATP binding. We describe the discovery of both an ATP- and a GTP-binding RNA aptamer in two independent selection experiments (both from poly-A starting sequences) and characterize their binding affinity and specificity.

## Results

In a typical SELEX (Systematic Evolution of Ligands by EXponential Enrichment) experiment the sequence diversity is high at the beginning and is progressively reduced during each selection step. In contrast, here we start with a maximally simple, single homopolymeric “seed” sequence and introduce sequence diversity before each selection round by error prone replication. Unlike the fully random sequence pool of a standard SELEX experiment, this creates a “cloud of genotypes” connected by mutation to a single “seed” sequence, akin to a quasispecies in viral replication. We chose to initiate selections from single poly-A and poly-U seed sequences as these more likely reflect the prebiotic scenario, as suggested by the increased yield and length distribution of poly-AU RNA polymers obtained by non-enzymatic non-templated polymerization compared to mixed sequences^11, 12^. These also avoid the complications arising from the propensity of poly-G, poly-C sequences to form highly stable non-canonical secondary structures (such as G4 quadruplexes and i-motifs) that hinder efficient replication.

Our experimental setup follows, with slight modifications, the canonical SELEX scheme for nucleotide aptamer selections^15^ (Figure 1). We carried out five independent selection experiments each starting from a single sequence with two sequences (T4, T6) with a poly-U segment and three sequences (T5, T7, T8) with a poly-A segment, each flanked by unique and different primer binding sites, to avoid cross contamination (Fig. 1, Supplementary Table 1). Furthermore, unlike previous selection experiments that use C8-linked ATP agarose, we used ATP agarose with the ATP linked via the gamma phosphate^16^. Our protocol includes an error-prone PCR step according to standard protocols (0.2 mM NTPs, 1 mM MnCl2)^17^ for every selection round including for the initial pre-selection repertoire (R0), to introduce sequence diversity. Deep sequencing T5 and T8 round 0 pools (T5R0, T8R0) (before selection) indicated a very high average mutation rate of up to 3 x 10^-1^, presumably due to a strong bias against amplification of the homopolymeric stretches and with a mutational bias (in T5R0) towards the 3’ end of the template (Supplementary Fig. 1). The challenges of replicating the unusual homopolymeric sequence may add additional bias and adaptive pressure for mutation away from homopolymeric runs, thus leading to elevated mutation rates in error prone PCR at least in the first round. Presumably due to the profound tendencies for slippage upon copying poly-dA or -dT stretches in PCR we also observed a strong tendency for truncation in sequence pools. We therefore included a gel-purification step into the selection cycle to enrich (near) full length sequences.

**Fig. 1.**
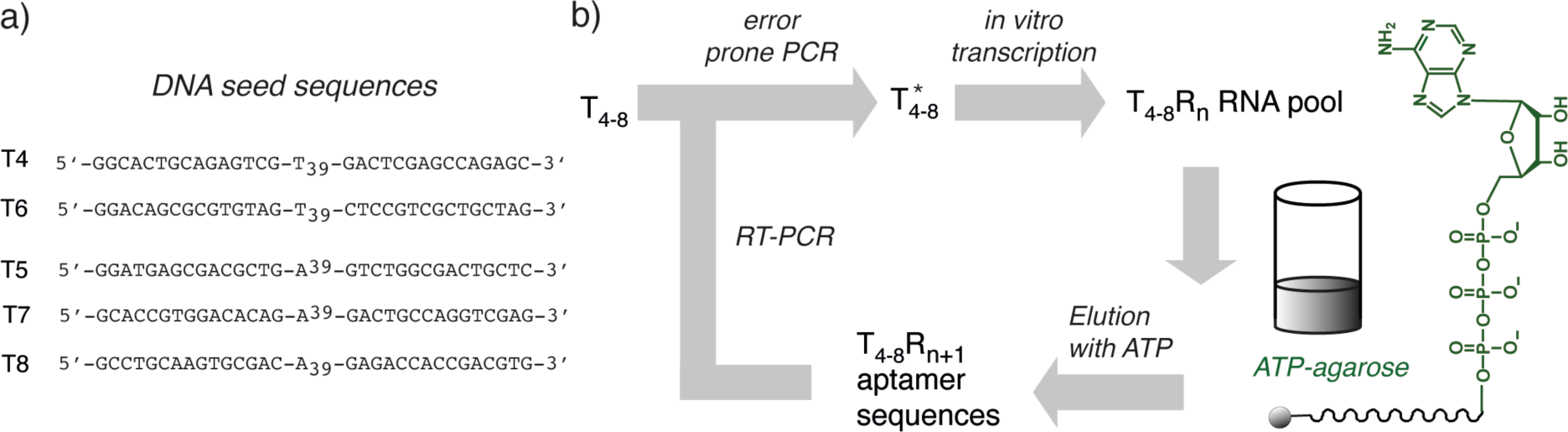
Schematic representation of the mutation selection experiment. a) Instead of a random sequence DNA library as for standard SELEX procedures, the experiment starts with a single poly-T (T4, T6) or poly-A (T5, T7, T8) sequence flanked by different primer binding sites. b) Principal workflow according to the standard SELEX protocol. T4-8 seed sequences are diversified by error prone PCR to yield sequence pool T*, which is transcribed, incubated with ATP agarose (γ-phosphate-linked), non-bound sequences (flow-through) are discarded and RNA sequences bound to the ATP matrix (ATP aptamers) are eluted with ATP. RNA sequences are reverse transcribed to cDNA and amplified by PCR (RT-PCR) and re-diversified by error prone PCR for another round of selection.

We performed 8 mutation-selection rounds from sequence T5, 16 rounds with sequences T7 and T8 (with increased selection stringency after round 12 (Supplementary Materials & Methods)) and 24 rounds with sequences T4 and T6.

Next, we searched all selections for appearance of the canonical RNA ATP aptamer loop motif^15^, which had previously been observed multiple times in unrelated selections^18–20^. Said motif comprises a conserved stem-loop-structure with an essential bulged G opposite the recognition loop (Figure 2a). Only 7 of the 11 nucleotides in the purine-rich recognition loop are conserved (GGNAGANNNTG), while even individual mutations of this conserved residues do not completely abolish binding. While the stem regions are essential, their composition has likely only minor influence on aptamer function, however at least one GC base pair in each stem is likely preferred for the formation of stable stems. We observed only sporadic appearance of the motif in the poly-U (T4, T6) selections even after 24 rounds of selection. In contrast, in poly-A (T5, T7, T8) selections we observed an increase of both the perfect and relaxed canonical (5’-GGAAGAAAATG-3’ (AAA) / 5’-GGNAGANNNTG-3’) (NNN) and (to a lesser extent) the reverse motif (5’-GTAAAAGAAGG-3’ (AAA_rev_) / 5’-GTNNNAGANGG- 3’)(NNN_rev_) in T5R8 (Fig. 2b) and the T7-, T8R12 pools (Supplementary Fig. 2), after which (in T7, T8) frequency dropped, possibly related to the more stringent selection conditions for the final 4 rounds of selection.

**Fig. 2.**
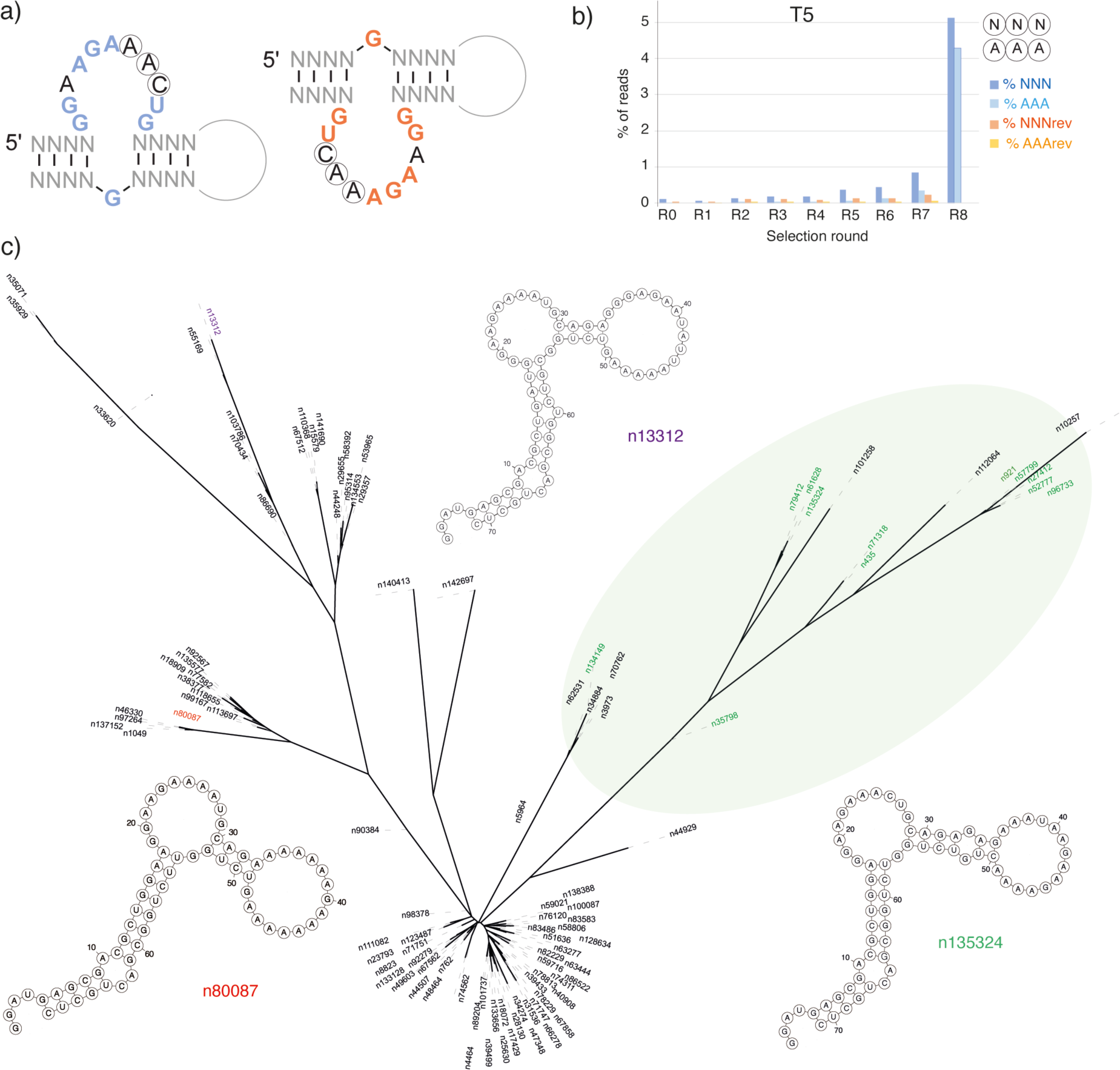
Emergence of ATP aptamer motifs. a) Canonical (left) and reverse ATP aptamer motif (right) with conserved bases in blue (canonical motif) and orange (reverse motif). NNN non-essential loop residues are circled. b) Ratio (%) from total unique sequences of the minimal (GGNAGA**NNN**TG (NNN) and near canonical GGAAGA**AAA**UG (AAA) ATP aptamer loop motif for every round (R0-R8) of the T5 selection experiment. c) Phylogenetic tree of the ATP aptamer motifs in T5R8 sequence pool (see Supplementary Fig. 3)(perfect match motifs in green) and predicted secondary structures of perfect match (n135324) and imperfect match (n13312, n80087) hits. Note the canonical ATP aptamer loop / G-bulge motif appears in all, with variations confined to the stem regions.

Encouraged by this we next searched for a perfect (or near perfect) match for the canonical ATP binding motif (including the bulged G)(leveraging the search algorithm of^21^). While no unambiguous hits were found in the T7R12 and T8R12 selections (despite appearance of the loop motif), in the T5R8 selection, we obtained >100 hits with divergent stem sequences (Fig. 2c, Supplementary Fig. 3) (of which 15 comprised a perfect match with the canonical motif). Several of these showed predicted canonical (or near canonical) ATP aptamer secondary structure motifs (as judged by RNAfold^22^).

As perfect matches to the canonical ATP aptamer motif had previously been shown to invariably bind to ATP^21, 23, 24^, we chose examples of more divergent but highly enriched sequences from both the T5 and the less promising T4, T6, T7 and T8 selections and compared their ATP binding activity to the starting seed sequences T4-T8 by ATP-agarose column elution. Among these we identified several motifs with putative ATP binding activity as judged by column elution profiles (T4R24/377, T5R8 / 359, T6R20 / 388, T7R16 / 401, T8R16/ 409) (Fig. 3a, Supplementary Fig. 4, Supplementary Table 2). However, among these only T5R8 / 359 and T8R16 / 409 showed strong ATP binding compared to their seed sequences and we focused further analysis on these two motifs.

**Fig. 3.**
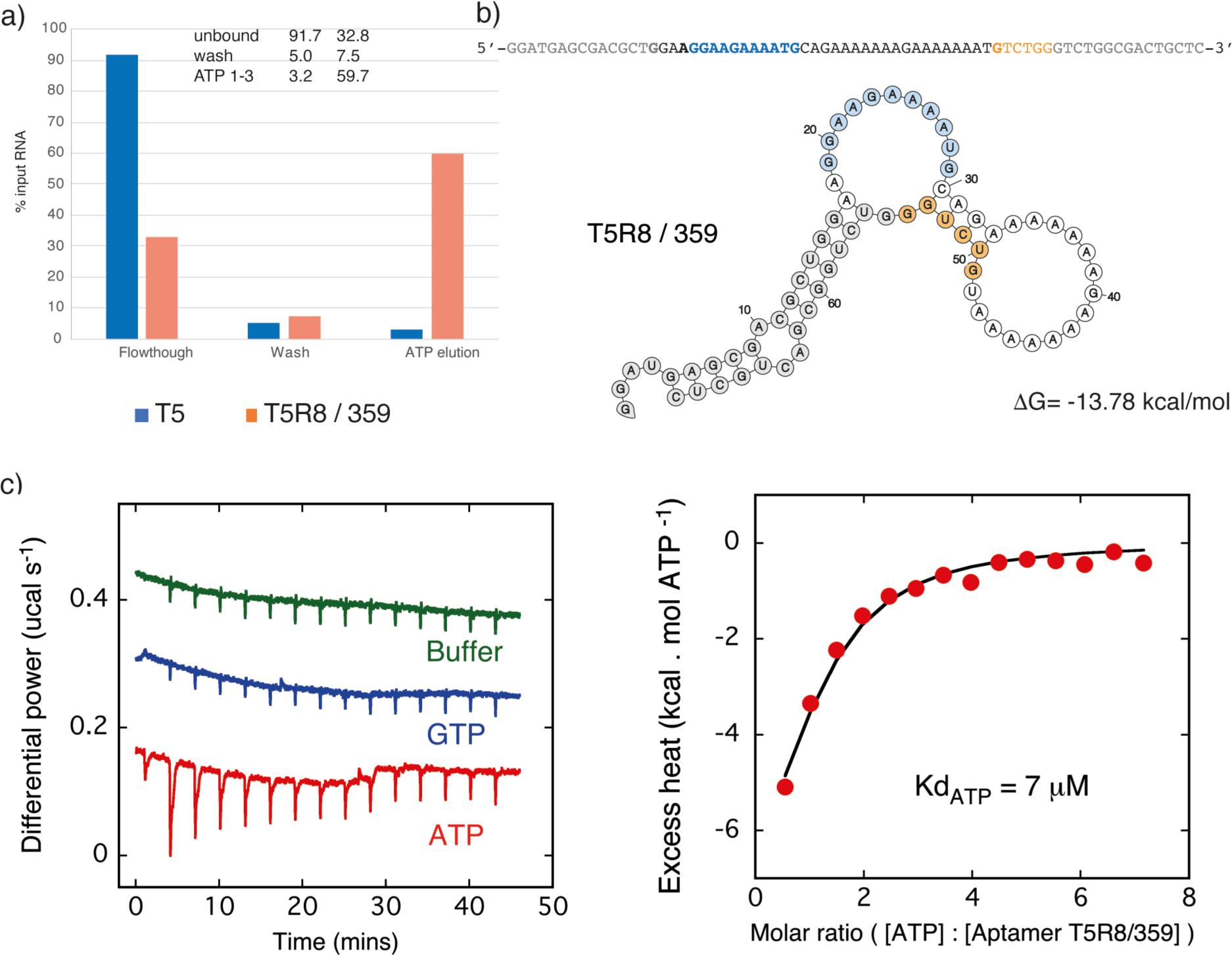
ATP Aptamer T5R8 / 359. a) ATP-agarose column elution of T5 seed sequence (blue) and the selected ATP aptamer T5R8 / 359 (orange). b) Sequence and predicted secondary structure. T5R8 / 359 aptamer with conserved primer sequences (grey), nucleotides corresponding to the canonical ATP aptamer loop motif (cyan) and nucleotides deriving from partial duplication of the primer sequence (orange) shown. c) ITC binding experiments : raw data showing additional binding heat for ATP titration compared to GTP or buffer controls (left panel) and integrated excess heat of ATP titration fit to a simple one site binding model (right panel)

T5R8 / 359 comprises a purine rich loop of 11 nucleotides flanked by two dsRNA regions and an essential G residue (bulged G) opposite the loop (Fig. 3b), thus differing by just 2 mutations from the canonical ATP aptamer motif. The essential opposing bulged G appears to originate from a partial (6 nt) duplication of the 3’ primer sequence (Figure 3b). In concordance with this T5R8 / 359 without the 6 nt 3’ primer insert (T5R8 / 359Δ6) showed no ATP binding as judged by column elution. (Supplementary Table 2). Despite an unpaired G.A next to the canonical G-bulge, T5R8 / 359 can adopt a canonical ATP aptamer-like fold, providing an interesting variation on the ATP binding motif. Where previously, a single G-bulge was considered essential for ATP binding^15, 23^, T5R8 / 359 suggests that an expanded loop and a G-G-bulge also support formation of a specific ATP binding pocket. This relaxes the search criteria for ATP aptamer motifs and suggest that there may be many more ATP-binding aptamers in the T5R8 pool than the >100 canonical motifs detected (Fig. 2c, Supplementary Fig. 3) and that more relaxed search criteria might be applied to identify ATP aptamer motifs in both SELEX experiments and in genomic RNA pools^21, 24^.

T8R16 / 409 from a different selection also showed clear ATP binding as judged by column elution (Fig. 4a, Supplementary Fig. 4) despite showing no sequence (or secondary structure) similarity with the canonical ATP aptamer motif. Notably, truncation by deletion of either (620) or both (T8R16 / 409core (105)) primer sequences resulted in no difference in ATP-binding as judged by column elution profiles, while the deletion of a central stretch of nucleotides abolished binding. While CTP and UTP elution profiles were similar to ATP, T8R16 / 409core appeared to be more readily eluted with GTP (Supplementary Fig. 5, Supplementary Table 3), pointing towards specificity for GTP.

**Fig. 4.**
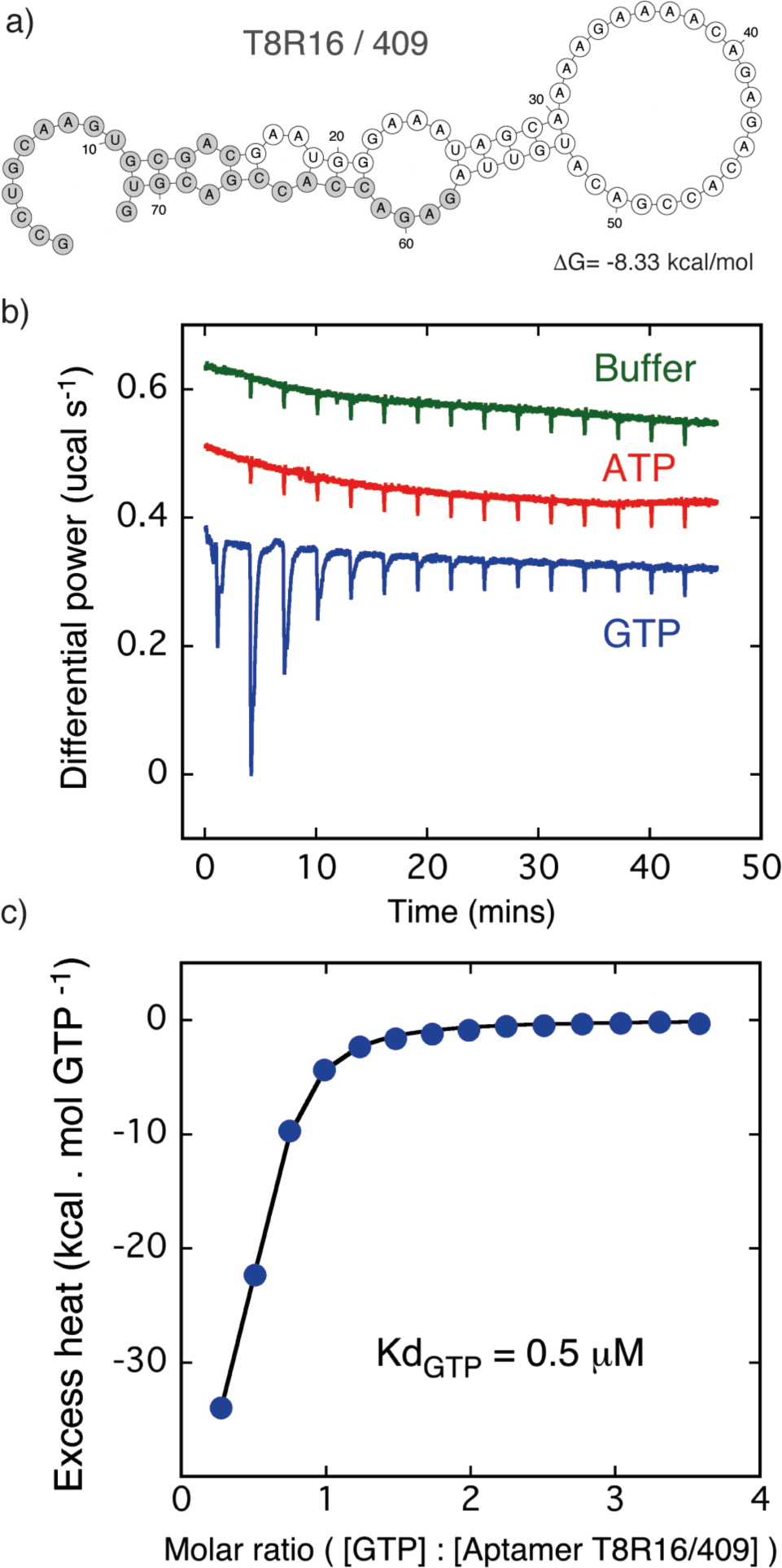
GTP Aptamer. a)Predicted secondary structure (RNA fold^22^) of T8R16 / 409 aptamer with conserved primer sequences (grey). b, c) ITC binding experiments. b) raw data showing additional binding heat for GTP titration compared to ATP or buffer controls and c) integrated excess heat of GTP titration fit to a simple one site binding model.

Next, we sought to measure specific binding affinities of these two most promising aptamer sequences T5R8 / 359 and T8R16 / 409 by isothermal titration calorimetry (ITC). Binding experiments were performed three times with similar results. The fitted isotherm indicated a Kd of aptamer T5R8 / 359 binding ATP of around 7 µM (Fig. 3c,d) while aptamer T8R16 / 409 binding GTP showed a more than 10x higher affinity with a Kd around 0.5 µM (Fig. 4b,c). Both showed remarkable selectivity with no binding detectable against the other nucleotide. Indeed the observed integrated heats were identical to those obtained by titrating the nucleotides into buffer alone suggesting no interaction at a level detectable in these experiments.

This is especially striking for the T8R16 /409 aptamer, which shows this strong selectivity against ATP, despite selection on an ATP-agarose resin as well as clear binding to and elution from the resin by ATP (as well as other ribonucleotides). As column elution assays can detect binding affinities in the mid to high µM range, this may simply reflect a promiscuous, but lower affinity binding of T8R16 / 409 to ATP (and indeed other nucleotides). The affinity of aptamer T8R16 / 409 for GTP yielded a reliable fit to a stoichiometry of around 0.5. The underestimate from an expected 1:1 value probably reflects potential errors in the concentrations of both aptamer and nucleotide, which were not easy to quantify, as well as the possibility of unstructured or misfolded aptamer material in the experiment. In light of the observed value for GTP we constrained the stoichiometry at this same value of 0.5 when fitting the ATP data. The issue of exact active concentration of materials used introduces some additional uncertainty into the precise values of the Kd constants measured. However, the experiments clearly demonstrate the binding of nucleotides by both these aptamers and their ability to discriminate between ATP and GTP in the µM and sub-µM regime of affinities.’

## Discussion

Here we have described the discovery of specific ATP- and a GTP-binding RNA aptamers by cycles of mutation and selection starting in each case from a single, non-functional, largely unstructured seed sequence (Fig. 1, 5, Supplementary Fig. 6). In two of five selection experiments we recover nucleotide binding aptamers with µM binding affinities for either ATP or GTP (as measured by ITC) (Figs 3, 4) as well as potentially hundreds of *bona fide* ATP aptamers as judged by the appearance of the canonical ATP aptamer motif (Fig. 2, Supplementary Fig. 3). Thus, the emergence of function is not dependent on highly diverse random sequence pools, but can arise from maximally simple, single sequences with minimal informational and molecular complexity with implications for the understanding of adaptive landscapes^8, 25^ and the potential emergence of function from non-functional areas of the genome (“junk” DNA)^26^.

**Fig. 5.**
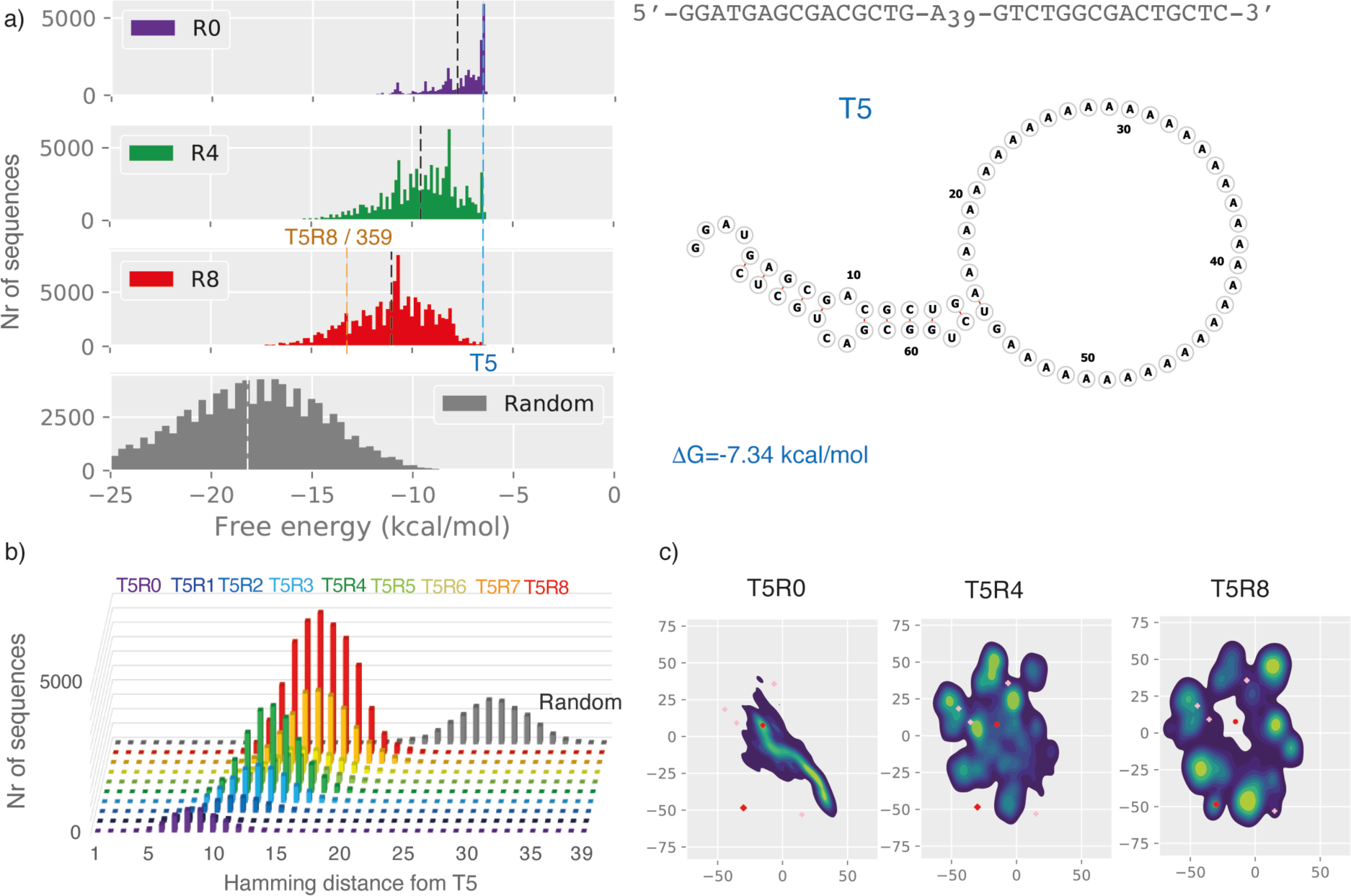
Sequence pool development during selection. a) (left panel) RNA folding energy distributions for T5R0 (purple), R4 (green), R8 (red) compared to random sequence pools (median: black / white dotted line). (right panel) Sequence, predicted secondary structure and folding energy (ΔG (cyan dotted line)) (RNA fold^22^) of T5 seed sequence. b) Hamming distance distribution of T5 sequence pool over the course of selection (R0-R8) with reference to a random sequence pool (grey). c) Sequence space evolution as shown by t-stochastic neighbour embedding (tSNE) for T5R0, R4 and R8 (T5 seed sequence (red dot), T5R8 / 359 aptamer (red diamond), ATP aptamer motif perfect hits (Fig. 2, pink crosses)(see Supplementary Fig. 8 for full trajectory).

The emergence of new function by iterative mutation and selection from a single (or very few) starting sequences has precedents in biology e.g. in virology (quasispecies)^27^, the avian immune system and hypermutating B-cell lines^28^. Examples also include model experiments in the ribozyme field morphing one ribozyme function into another^29^. However, these examples differ from our work in that new function does not emerge *de novo.* but from already stably folded (and functional) protein (or RNA) domains. In fact we consciously chose starting sequences with minimal structural or informational complexity (Fig. 5, Supplementary Fig. 6) comprising a central homopolymeric stretch of A39 or T39 (U39), flanked by conserved 15 nt primer sequences for amplification and subjected them to iterative rounds of diversification by random mutation and selection and analysis. Sequencing analysis suggested very high mutation rates (Supplementary Fig. 1) and therefore rapid diversification of the seed sequences radiating out from a single point in sequence space (Supplementary Figs 7-9), presumably due to the difficulties in accurately copying the mononucleotide repeat sequence. However, even though the sequence and hence molecular diversity of the pools in all selections continued to expand during the selection process (as judged by Hamming distance), they never reached the complexity of a fully random pool and thus maintain a significant sequence bias and imprint of the seed sequence even after 24 rounds of mutation / selection (Fig. 5, Supplementary Fig. 7). Nevertheless, new sequence features continue to emerge during the course of selection as shown by principal component analysis by t-stochastic neighbor embedding (tSNE)^30^ (a two-dimensional projection visualization of sequence space evolution) (Fig. 5, Supplementary Figs. 8, 9). This shows that even from such biased sequence pools, functional sequences can emerge as we demonstrate

While high mutation rates lead to rapid diversification, they also hinder the fixation and stable inheritance of a given phenotype. Furthermore, they might bias selection not just towards small motifs^31^, but also towards RNA folds that are both abundant and robust (stable) (such as hairpin and hairpin-like motifs)^32, 33^ that can withstand high mutation rates while maintaining overall structure and function even if they are not the most “fit”, i.e. best adaptive solutions, in a process which has been termed “survival of the flattest”^34^. Indeed, in all cases overall selected sequence pools become more structured and more stably folded (as judged by secondary structure prediction) than starting sequences (Supplementary Figs 4, 6). However, in none of the selections do they reach the median stability of a random pool and this trend may therefore simply reflect increasing sequence diversification. Although all selections yield stably folded motifs with detectable, if weak, ATP binding, as judged by column elution (Supplementary Fig. 4, Supplementary Table 2), the folding stabilities of the highly functional aptamer binding motifs (T5R8 / 359, T8R16 / 409) are at higher (359) respectively lower (409) end of median stability for their rounds (Fig. 5a, Supplementary Fig. 6). Clearly, while structural pre-organization can enhance the success of aptamer discovery^9^, and may buffer a phenotype against high mutation rates, the extent of folding and structural stabilization does not appear to correlate with function at the global level.

This canonical ATP aptamer fold appears to be a privileged molecular solution for ATP binding, as it has been isolated multiple times by *in vitro* selection on ATP-agarose in independent selections against ATP containing cofactors such as NAD^18^ and SAM^19, 35^ and was even identified in several bacterial and eukaryotic genomes^21, 24^. This parallels the case of the Hammerhead ribozyme motif^36, 37^ and suggests that both of these motifs may represent minimal optimal solutions in RNA sequence space. The isolation of the canonical ATP aptamer motif in the T5 selection may also indicate its resilience to high mutation rates.

The isolation of a GTP binding aptamer (T8R16 / 409)(Fig. 4) from one of the ATP selection experiments may seem fortuitous, but is less surprising considering the previous discovery of numerous GTP binding aptamers at a relatively short mutational distance from the canonical ATP aptamer motif^38^. Furthermore, GTP aptamers in general appear to be much more structural and functional diverse and may therefore also be more abundant in sequence space. Indeed, “CA” repeats as short as 3 nucleotides^39^ or G-quadruplex motifs^40^ have been reported to bind GTP. Finally, while the GTP binding aptamer only binds GTP in solution (as judged by ITC), it clearly binds to the ATP-agarose resin (both as the full-length T8R16 / 409 aptamer as well as the aptamer core (T8R16 / 409core)), from which it is slowly eluted by ATP, CTP, UTP and more efficiently by GTP (Supplementary Figs 4, 5, Supplementary Table 3). This discrepancy is likely to reflect the much higher sensitivity of the column elution assay, which detects much lower (high µM to mM) affinities that are outside the range of ITC experiments designed to quantify sub-µM affinities.

How structure and function (the phenotype) of a biopolymer relates to its sequence (the genotype) is often discussed in terms of a fitness landscape of peaks and valleys with evolution viewed as a random walk in the direction of adaptive peaks. The shape e.g. degree of “ruggedness” of such landscapes is an active area of research^8, 25, 41, 42^ and has clear implications for both the likelihood of reaching optimal fitness peaks and the best adaptive strategies. For example, comprehensive studies of both a GTP aptamer and a kinase ribozyme fitness landscape found them to be substantially rugged with few neutral networks (i.e. mutational paths) connecting adaptive peaks^8, 25^. However, the discovery of nucleotide binding aptamers from single starting sequences as described here suggests that there must be permissive mutational trajectories connecting adaptive peaks even to single, unstructured, non-functional sequences. While it seems that such trajectories may be difficult to reconcile with the substantial ruggedness of empirical fitness landscapes^8, 25^ (and the scarcity of neutral networks (adaptive paths connecting different (at least partially) functional points in sequence space via single mutational steps), the sequence of the selected T5R8 / 359 aptamer suggests that diversification by mutation alone may be insufficient and recombination may be an important route to access such trajectories (Fig. 3).

Our results underline the importance of context and contingency as the same poly-A core evolved very differently depending on the conserved flanking sequences (which are not mutated during selections). While the canonical purine-rich loop motif was enriched in all poly- A selections (T5, T7, T8) (Fig.2, Supplementary Fig. 2), the key opposing G-bulge required for high affinity ATP binding motif only arose in the T5 selection (Supplementary Fig. 3) arising from a partial duplication of the 3’ flanking sequence. Furthermore, the variable success of aptamer discovery may reflect the availability of accessible vs. inaccessible adaptive trajectories e.g. through formation of favorable RNA secondary structures^43, 44^ or the respective mutational distance of the seed from adaptive peaks. Indeed, the canonical purine-rich ATP binding motif^15^ is more readily accessible from a poly-A than a poly-U sequence, both with regards to mutational distance and the increased likelihood of A > G transition vs. U > A/G transversion mutations. Interestingly, this dichotomy is a reflection of the SELEX process comprising transition between RNA and DNA intermediates (through reverse transcription into cDNA, PCR amplification followed by in vitro transcription), whereby only one RNA strand is available for selection. In a prebiotic scenario, where RNA would be replicated from RNA templates (either non-enzymatically or by RNA polymerase ribozymes)^4^, both RNA strands would potentially be available for selection and both poly-A- and poly-U -rich seed sequences would therefore be comparable starting points for evolution.

In biology, with the exception of specialized tissues like the immune system^28^ and some viruses^27^, genetic information is replicated with high fidelity and diversity derives from drift (a slow accumulation of nearly neutral mutations) and recombination. In contrast, current model systems of prebiotic RNA replication suggest an error prone process both for non-enzymatic RNA replication^45, 46^ as well as enzymatic replication by polymerase ribozymes^47^ combined with an innate tendency of RNA oligomers for recombination^48^. Our approach mimics such a scenario and indicates that highly error-prone replication (even beyond the Eigen error threshold^49^) does not only not impede the emergence of simple functional motifs, but may aid or even be required for the emergence of function from the biased prebiotic sequence pools or indeed single seed sequences.

## Acknowledgements

This work was supported by a postdoctoral fellowship no. 293387 from the Simons Foundation (FW), by the Medical Research Council (MRC) program grant program no. MC_U105178804 (PH, CJ) and a research collaboration between AstraZeneca UK Limited and the Medical Research Council-MRC-AstraZeneca Blue Sky Grant (BTP).

Correspondence and requests for materials should be addressed to P.H.

## Author contributions

FW and PH conceived and designed experiments. FW performed all experiments except ITC analysis of aptamer binding (CJ) and data analysis (with BTP). All authors analyzed data, discussed results and co-wrote the manuscript.

## Competing interests

The authors declare no competing interest.

## Data availability

The data that support the findings of this study are available from the corresponding author upon reasonable request.

## Code availability

Custom scripts used for analysis in this study can be found at GitHub:

## Materials and Methods

### Oligonucleotides

DNA and RNA oligonucleotides (listed in Supplementary Table 1) were from Integrated DNA Technologies and if necessary were purified, by denaturing PAGE (8M urea, TBE). The 2 poly-T (T2, T4) and 3 poly-A (T5, T7, T8) starting sequences consist of 69 nt, containing an either 39 nt poly-A or poly-T sequence flanked by different 15 nucleotide (nt) primer binding sites on both ends (Supplementary Table 1). The sequences were designed with non-overlapping primer binding sites to avoid cross contamination from different evolution-selection experiments.

### Evolution-selection experiments

The poly-A / poly-T DNA sequences (T2, T4, T5, T6, T7, T8) were PCR amplified (MJ Research Tetrad PTC-225 Thermal Cycler) (25 cycles), using the OneTaq hot start master mix (Promega) and corresponding primers (Supplementary Table 1), with the forward primer including the T7 RNA polymerase promoter sequence. PCR products were gel purified by agarose gel electrophoresis and isolated applying QIAquick columns (Qiagen), followed by in-vitro transcription applying the Megashortscript kit (Ambion). After DNase I (Ambion) treatment (30 min, 37°C), the RNA was isolated from the reaction mixture using the RNeasy kit (Qiagen) and the obtained crude RNA was purified by denaturing PAGE (8 M Urea, TBE).

The RNA sequences were first subjected to a negative selection step on sepharose, before selecting against ATP agarose. For the selection step 10 pmol of purified RNA was heated in reaction buffer (300 mM NaCl, 5 mM MgCl2, 20 mM TRIS pH 7.4) to 65°C for 5 min and cooled down at rt for 20 min. The RNA was added to 80 µl of sepharose (Sigma), that was prewashed with reaction buffer (3x 160 µl), in Costar microfilter spin columns (0.45µm, CA membrane, Corning), and was incubated for 1h at room temperature (rt). The flow-through was subsequently added to 40 µl of reaction buffer prewashed (3x 120 µl) ATP agarose (γ-phosphate-linked, 8-12µmol/ml, Innova Biosciences) in Costar microfilter spin columns and incubated for 1h at room temperature. The flow-through was discarded and the ATP agarose resin was washed three times with 160 µl of reaction buffer, followed by the addition of 120 µl of 5mM ATP (Roche) in reaction buffer and incubated for 30 min.

For the selection rounds 12-16 (T7, T8) and 20-24 (T2, T4) the columns were first eluted with 1 mM ATP in reaction buffer, followed by the 5 mM ATP elution. The flow-through was desalted using Vivaspin 500 (3000 MWCO) concentrators (Sartorius). The obtained RNA was reversed-transcribed and PCR-amplified using the Superscript III One Step RT-PCR system (Invitrogen) followed by an additional error-prone PCR step (20 cycles) using standard Taq DNA polymerase (HT Biotechnology) with equimolar concentrations of dNTPs (200 µM) but with the supplement of 1 mM MnCl2 (final conc.). The obtained DNA was purified and in-vitro transcribed to RNA as described above and the purified RNA was added into the next selection cycle. The sequences defined as round 0 corresponds to one round of standard PCR (25 cycles) and one round of error-prone PCR (20 cycles).

### Deep Sequencing

DNA products from different selection rounds were PCR amplified using extended primers including adaptor sequences for Illumina sequencing (Supplementary Table 1). The DNA was purified by agarose gel electrophoresis and extracted using the Qiagen gel extraction kit (Qiagen), followed by quantification by q-PCR (Brilliant III Ultra-Fast SYBR® QPCR, Agilent). Sequencing was performed on a MiSeq Illumina platform and sequencing data were processed by trimming of adaptor sequences, quality filtering, conversion to FASTA format and collapsing of sequences according to abundance using the Galaxy platform web server: https://usegalaxy.org. The occurrence of the ATP aptamer loop motif 5’-GGAAGAAAATG-3’ / 5’-GGNAGANNNTG-3’ / 3’-GTAAAAGAAGG-5’ (rev AAA) or 3’-GTNNNAGANGG-5’ (rev NNN) was analysed using FIMO (Find Individual Motif Occurrences) http://meme-suite.org/tools/fimo. RNA secondary structure prediction and display was carried using the ViennaRNA package 2.0^1^, specifically RNAfold either with or without aptamer loop motif contraints.

### Mutation rate analysis

We measured the mutation rate during error prone PCR (epPCR) through deep sequencing of round 0 (R0), which is the unselected output from epPCR of a poly-A stretch of 39 nucleotides. The mutation rate for each position was calculated with equation 1.

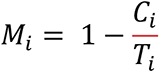

Where C_i_ is the number of correct nucleotides (A) at position i, and Ti is the total number of nucleotides at position i. The average mutation rate is the mean rate of all 39 nucleotides.

### Hamming distance analysis

When comparing converging or diverging sequence populations against a target sequence, we used the Hamming distance. Briefly, we computed this as the sum of positional mismatches for a given sequence, against a reference of the same length.

### Sequence space projections

To visualise sequence space of our aptamer selection experiments, we chose to reduce the dimensionality and project it into two dimensions using t-Stochastic Neighbour Embedding (t-SNE)^2^ implemented in the openTSNE package v.0.4.0^3^. Here, we used a perplexity of 50, a cosine metric, and principal component analysis for initialisation, with all other parameters as default. We used sequencing reads that were trimmed to contain the full 5 ’and 3 ’primer sequences, one-hot encoded into a binary triplet encoding, and padded with zeros out to 50 nucleotides.

For the projection figures, we merged and collapsed reads across all rounds, fit the t-SNE model as described above, and transformed the reads for each round to draw a mixed scatter and kernel density plot using matplotlib^4^ and seaborn (https://doi.org/10.5281/zenodo.592845). Canonical ATP aptamer motif hits were identified using RNABOB, written by Sean Eddy (Washington University School of Medicine, St Louis, MO; http://www.genetics.wustl.edu/eddy/software/#rnabob), and the following descriptor files, transformed using the fitted model and plotted as pink plus signs.

\# Canonical ATP aptamer motif strict

h1 s1 h2 s2 h2’ s3 h1’

h1 0:0 NNNN:NNNN

h2 0:0 NNNN:NNNN

s1 0 GGAAGAAACTG

s2 0 N[30]N

s3 0 G

\# Canonical ATP aptamer motif relaxed h1 s1 h2 s2 h2’ s3 h1’

h1 0:0 NNNN:NNNN

h2 0:0 NNNN:NNNN

s1 0 GGAAGAAANTG

s2 0 N[30]N

s3 0 G

### Phylogenetic tree generation

Canonical ATP aptamer motifs were identified using RNABOB as described above and a multiple sequence alignment performed with muscle v3.8.1551 using default settings^5^. With the aligned motif sequences (including 5 ’and 3 ’primer sites), a phylogenetic tree was calculated with MrBayes 3.2.7 single default parameters^6^. Phylogenetic trees (Fig. 2, Supplementary Fig. 3) were displayed using the EMBL iTOL online tool for the display, annotation and management of phylogenetic trees: (https://itol.embl.de/).

### Column and elution binding assay

RNA Sequences for the column binding assay, were generated by in vitro transcription from DNA templates generated by PCR amplification or by fill-in with primer T7X (Supplementary Table 1) using GoTaq (Promega). DNA was purified using QiaQuick (Qiagen), transcribed using the Megashortscript kit (Ambion) and purified using RNeasy (Qiagen). Cy5 labelled RNA was ordered purified (HPLC) from IDT. 40 µl ATP agarose was added to spin columns (0.45 µm, CA membrane, Corning) and equilibrated through washing with 3 x 120 µl buffer (300 mM NaCl, 5 mM MgCl2, 20 mM TRIS pH 7.4). 4 pmol RNA oligonucleotide in 80 µl buffer was heated to 65°C, cooled down for 15 min, added to the column and incubated for 1h at rt. After washing with 80 µl buffer, three to five subsequent ATP elutions (80 µl of 5 mM ATP in buffer) were performed, with a delay time of 5 min between each elution and the volume of the different fractions was reduced to ∼ 15 µl by speed-vac. After the addition of 30 µl of 8M urea/ 0.1 M EDTA the fractions were added to a 8M urea PAGE gel and were run at constant power (30 W). The gel was stained by SYBR gold, analysed by a Typhoon scanner and quantified using ImageQuant (GE Healthcare). Binding assays were repeated 2-3 times for each RNA sequence.

### Isothermal titration calorimetry

ITC experiments were performed at 25°C in 50 mM HEPES, 100 mM NaCl, 5 mM MgCl2, pH 7.6 buffer using a Malvern Panalytical both manual and auto iTC200 instruments. The RNA aptamers were loaded into the ITC cell at 6uM and the nucleotides titrated from the syringe at 100-200uM depending on affinity. Titrations were performed with 14 injections of 2.8 uL of nucleotide preceded by a small 0.5 uL pre injection that was not used during analysis. After baseline fitting and integration of excess heats the data were corrected using control heats observed for identical titrations of nucleotide solution into buffer. These control heats were small and similar to values obtained for buffer blank titrations. Corrected titrations were fit using Malvern Panalytical PEAQ software using a simple one set of sites binding model.

## Supplementary Figures

**Supplementary Fig. 1.**
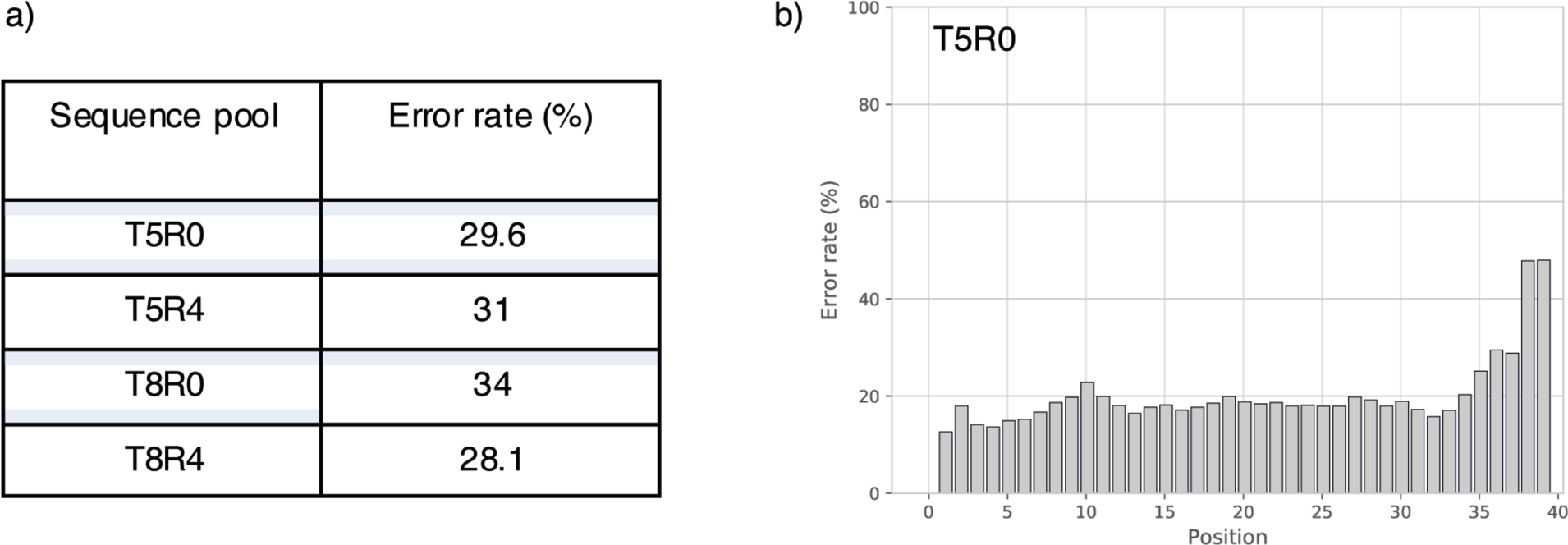
Mutation rates. a) Estimation of error rates of initial error prone PCR step for T5 and T8. The error rate was determined from nucleotide position 16-55 (1-39) (excluding the primer binding sites) applying the starting sequence (T5, T8) as reference. b) Estimation of positional error rates of initial error prone PCR step for T5R0 using seed sequence T5 as a reference. Note the increase of the error rate towards the 3’ end of the segment.

**Supplementary Fig. 2.**
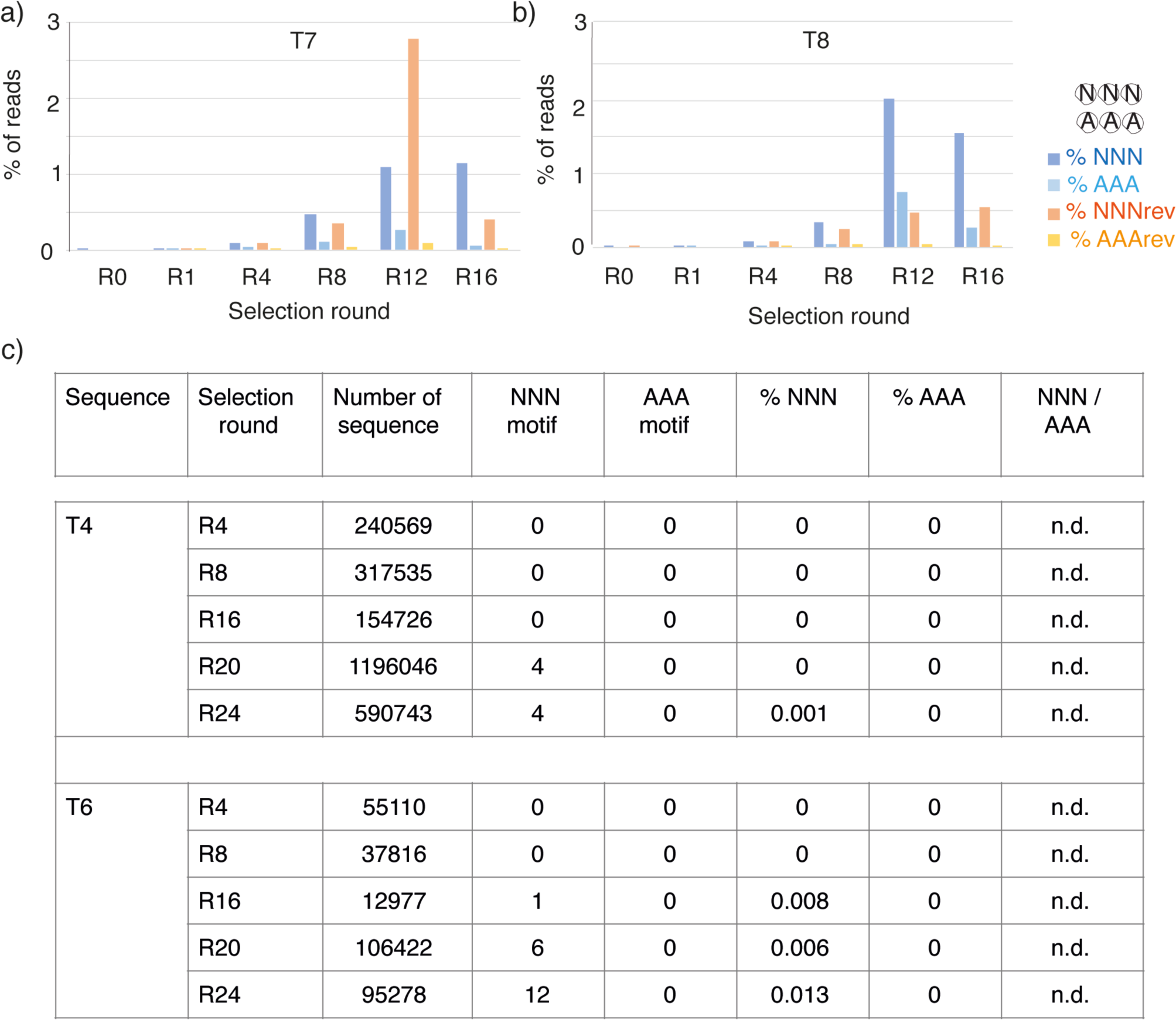
Appearance of aptamer motif loops (see Fig. 2a). a, b) Ratio (%) from total unique sequences of the minimal (GGNAGA**NNN**TG (NNN) and near canonical GGAAGA**AAA**UG (AAA) ATP aptamer loop sequence motif (Canonical ATP aptamer motif (blue, cyan) and reverse motif (red, orange) for selection rounds (R0, R1, R4, R8, R12, R16) of the T7 and T8 selection experiments. c) Sporadic appearance of canonical aptamer motif loops in the T4 and T6 selection experiments. Reverse loops were not detected.

**Supplementary Fig. 3.**
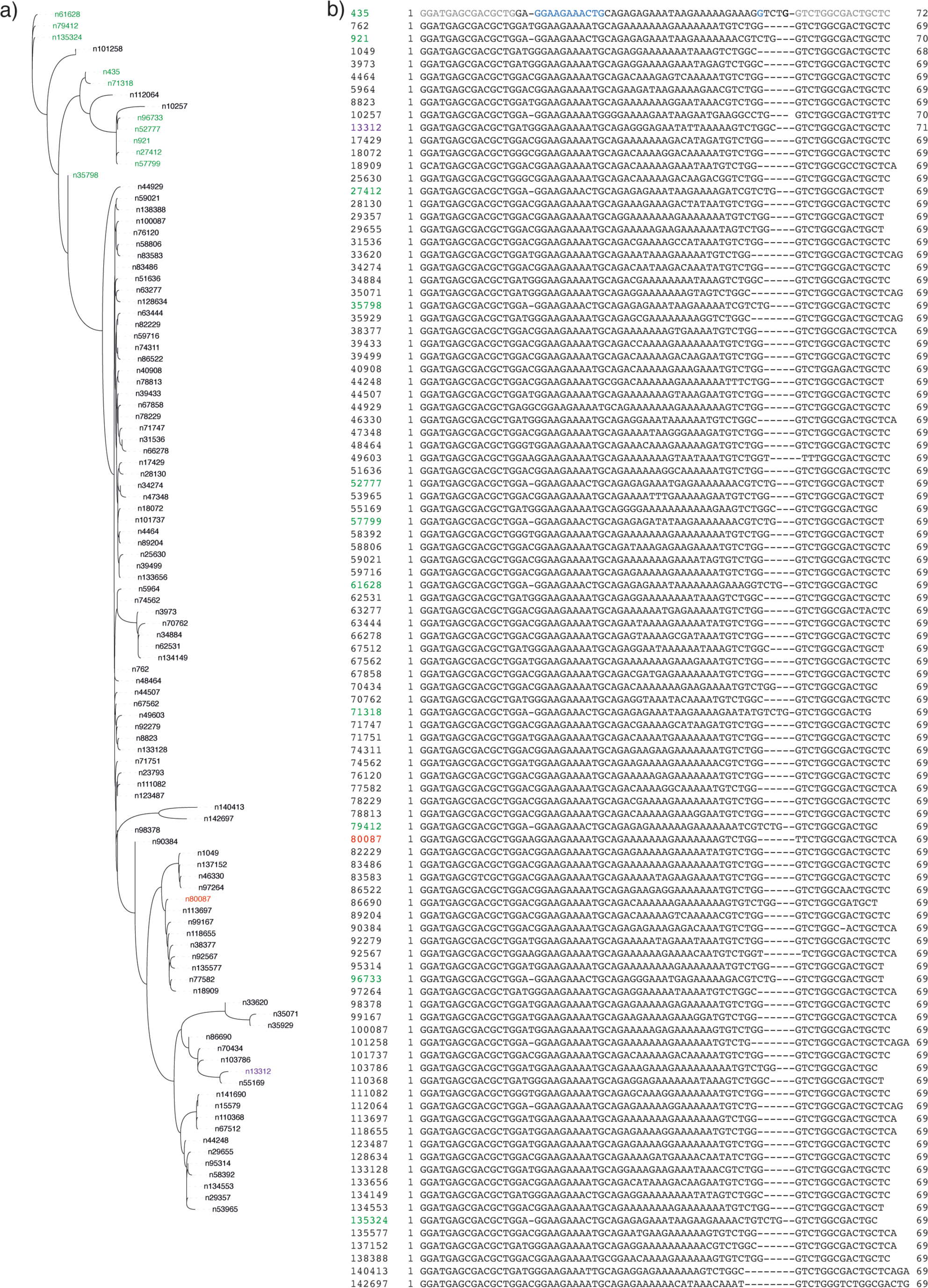
Aptamer motifs. a) Linear phylogenetic tree of b) predicted ATP aptamer motifs in T5R8 sequence pool (see Fig. 2). Primer sequences (grey), aptamer loop and G-bulge (blue).

**Supplementary Fig. 4.**
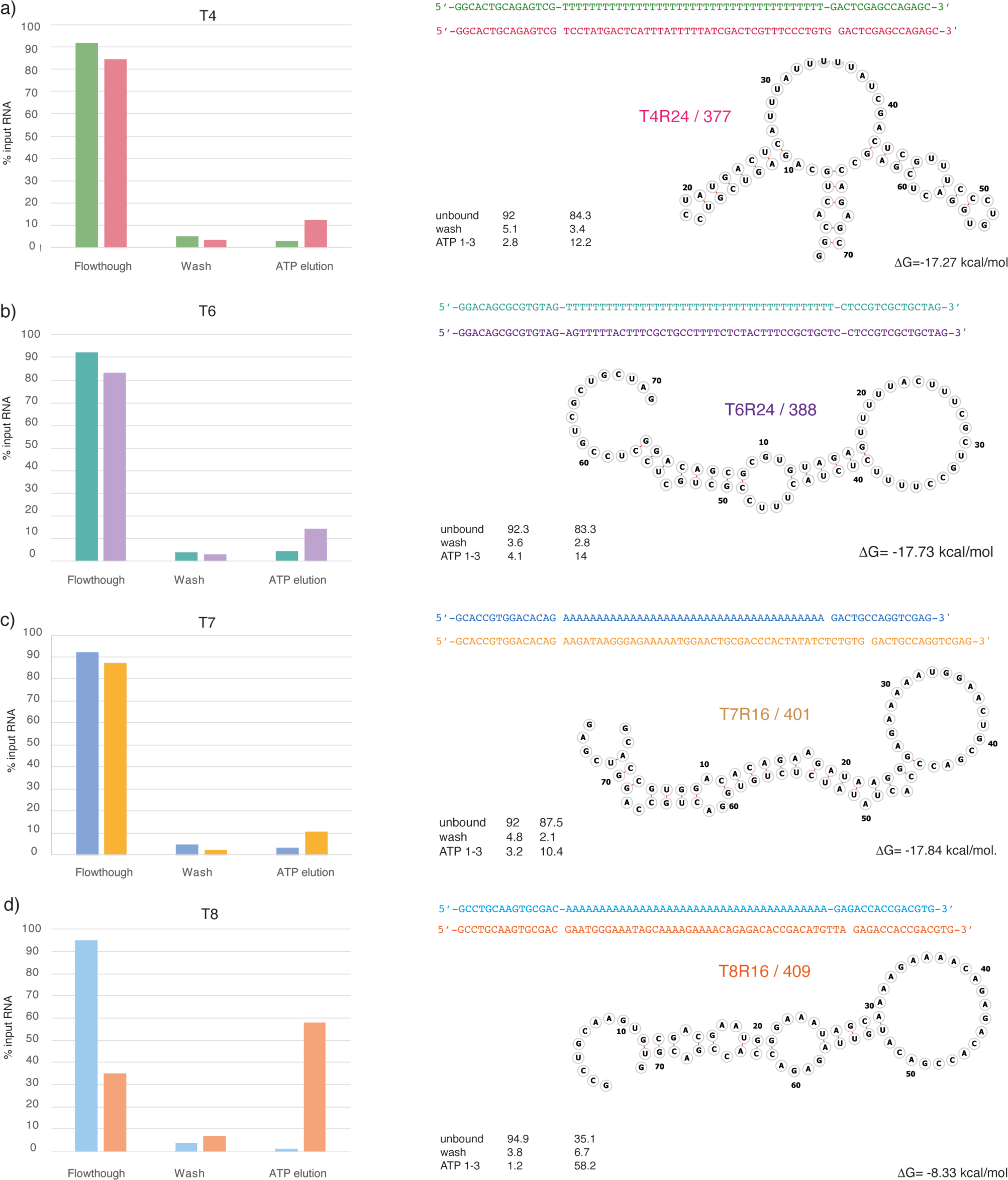
Putative ATP aptamer motifs. (Left panel) ATP-sepharose column elution of a) T4, b) T6, c) T7, d) T8 seed sequences and the selected putative ATP aptamers a) T4R24/377, b) T6R24/388, c) T7R16/401, d) T8R16/409. (Right panel) Sequence, predicted secondary structures and stabilities (ΔG)(as judged by RNAfold^1^) of putative aptamers.

**Supplementary Fig. 5.**
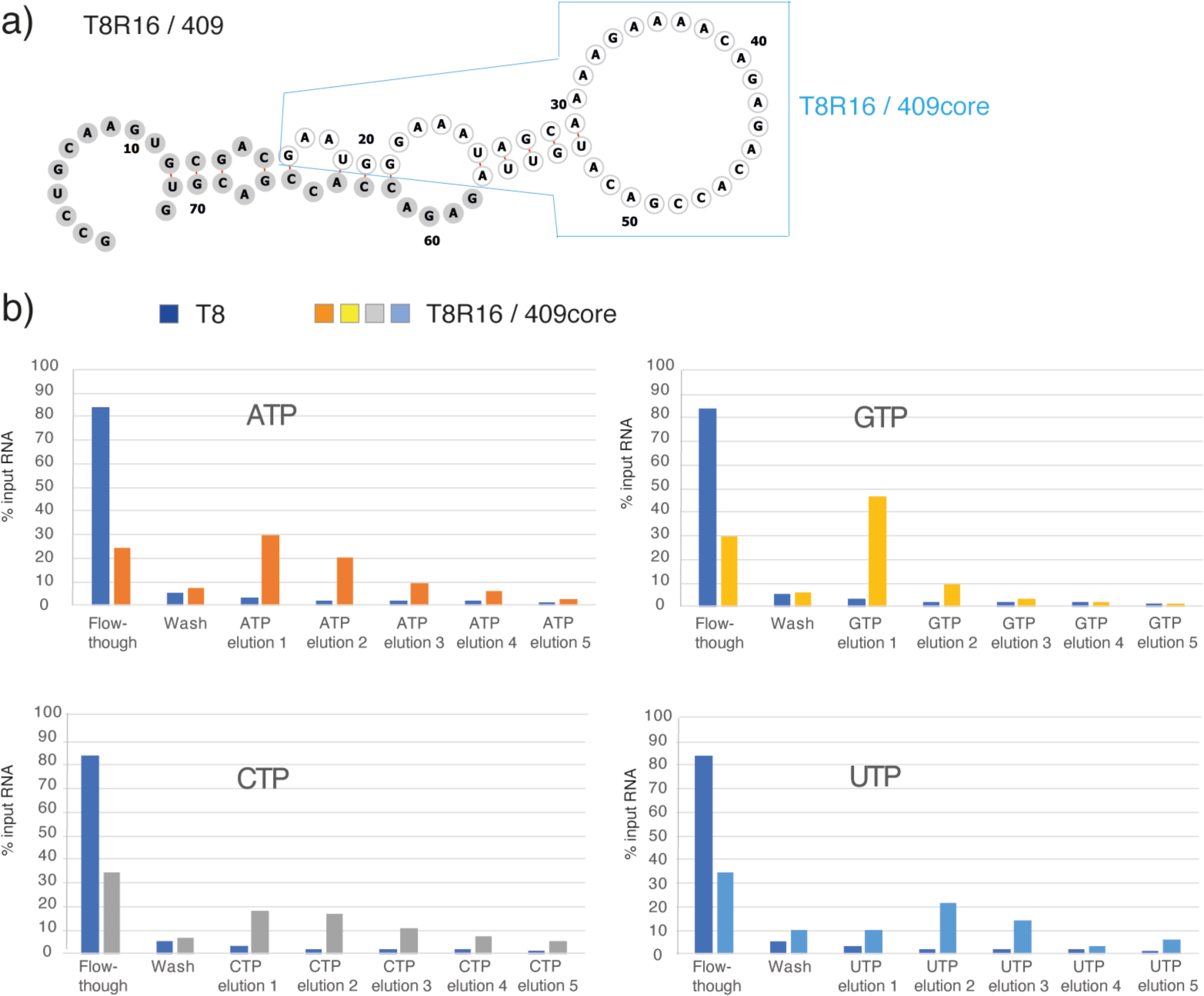
Putative nucleotide binding aptamer T8R16/409. a) Predicted secondary structure (RNAfold^1^) of T8R16/409 aptamer with conserved primer sequences (grey) and core domain (T8R16/409core) (cyan box).b) ATP-agarose column elution of T8 seed sequence (blue) and T8R16/409core with ATP (orange), GTP (yellow), CTP (grey) and UTP (cyan) (see Supplementary Table 3). Note that T8R16/409core is eluted by all 4 nucleotides, but most efficiently by GTP.

**Supplementary Fig. 6.**
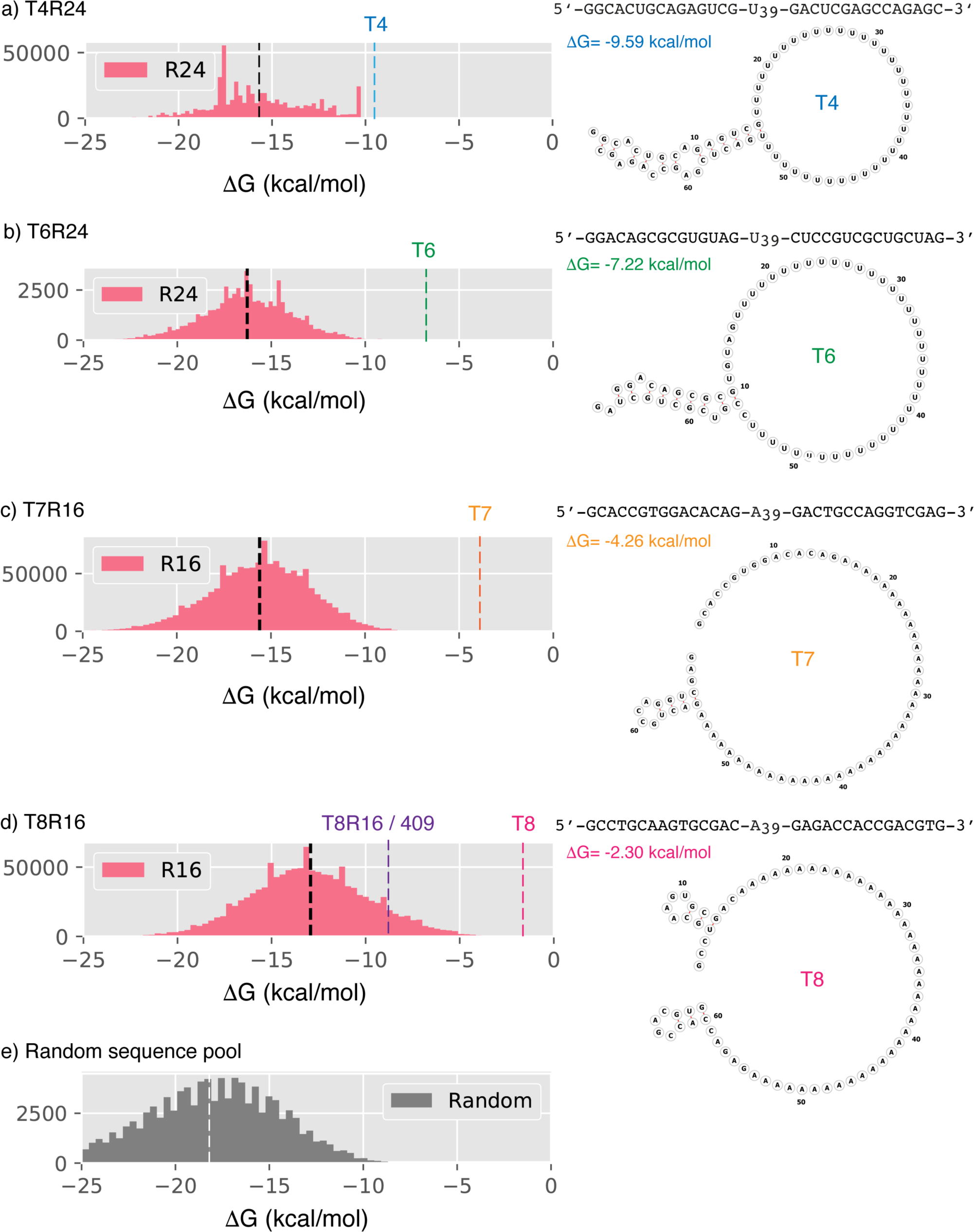
RNA folding. (Left panel) RNA folding energy distributions for a) T4R24, b) T6R24, c) T7R16, d) R8R16, e) random sequence pools (median: black ((e) white)) dotted line. (right panel) Sequence, predicted secondary structure and folding energy (ΔG (cyan dotted line)) (RNA fold^1^) of T4, T6, T7, T8 seed sequences.

**Supplementary Fig. 7.**
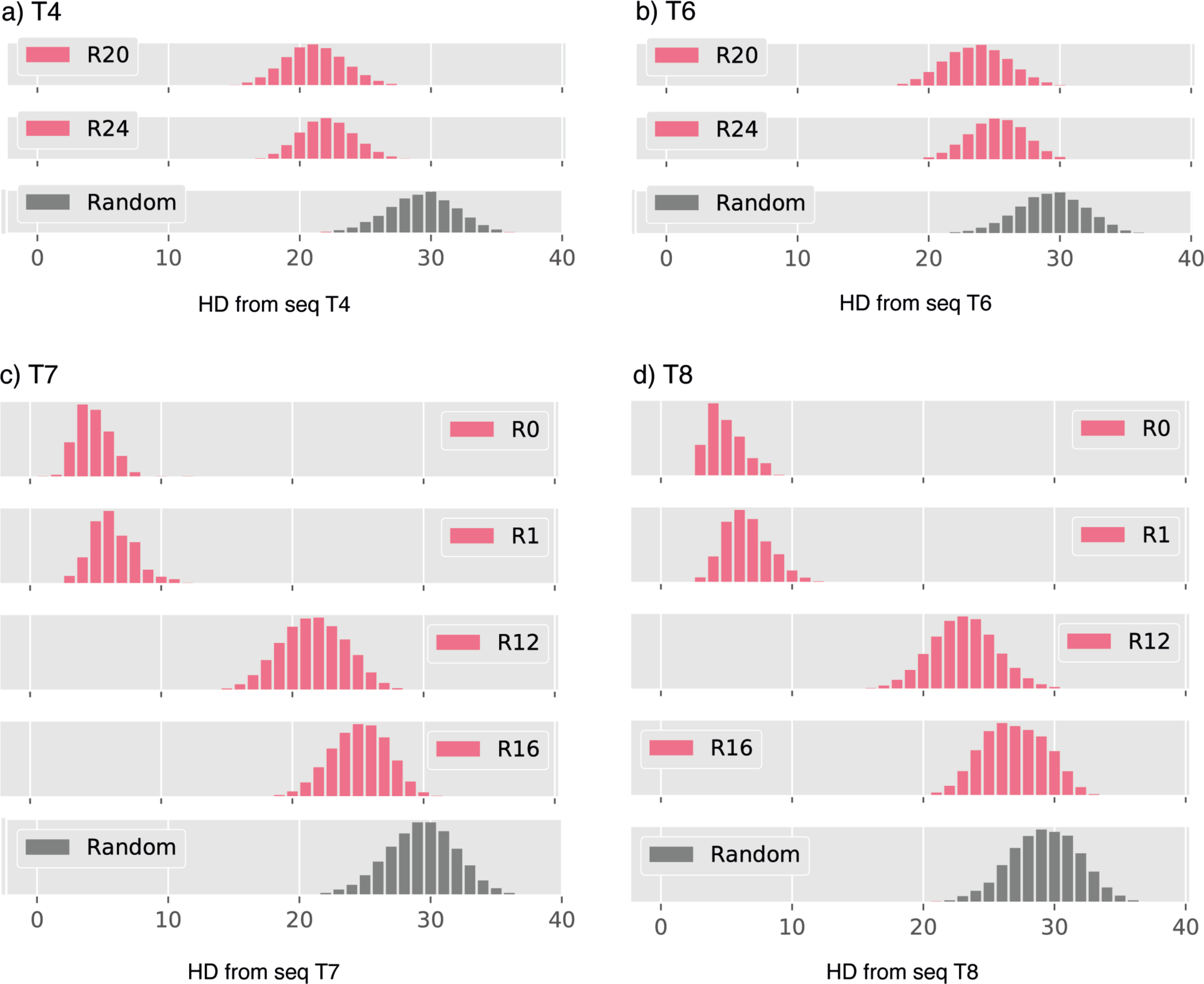
Hamming distance (HD). Hamming distance distribution of a) T4, b) T6, c) T7, d) T8 sequence pools at different points of the selection (R0-R24) with reference to a random sequence pool (dark grey). A Hamming distance of x between two sequences 1 & 2 means that x mutations are required to change sequence 1 into sequence 2. Therefore, random sequences (grey) will (on average) be at a mutational (HD) distance of 29-30 from any homopolymeric sequence.

**Supplementary Fig. 8.**
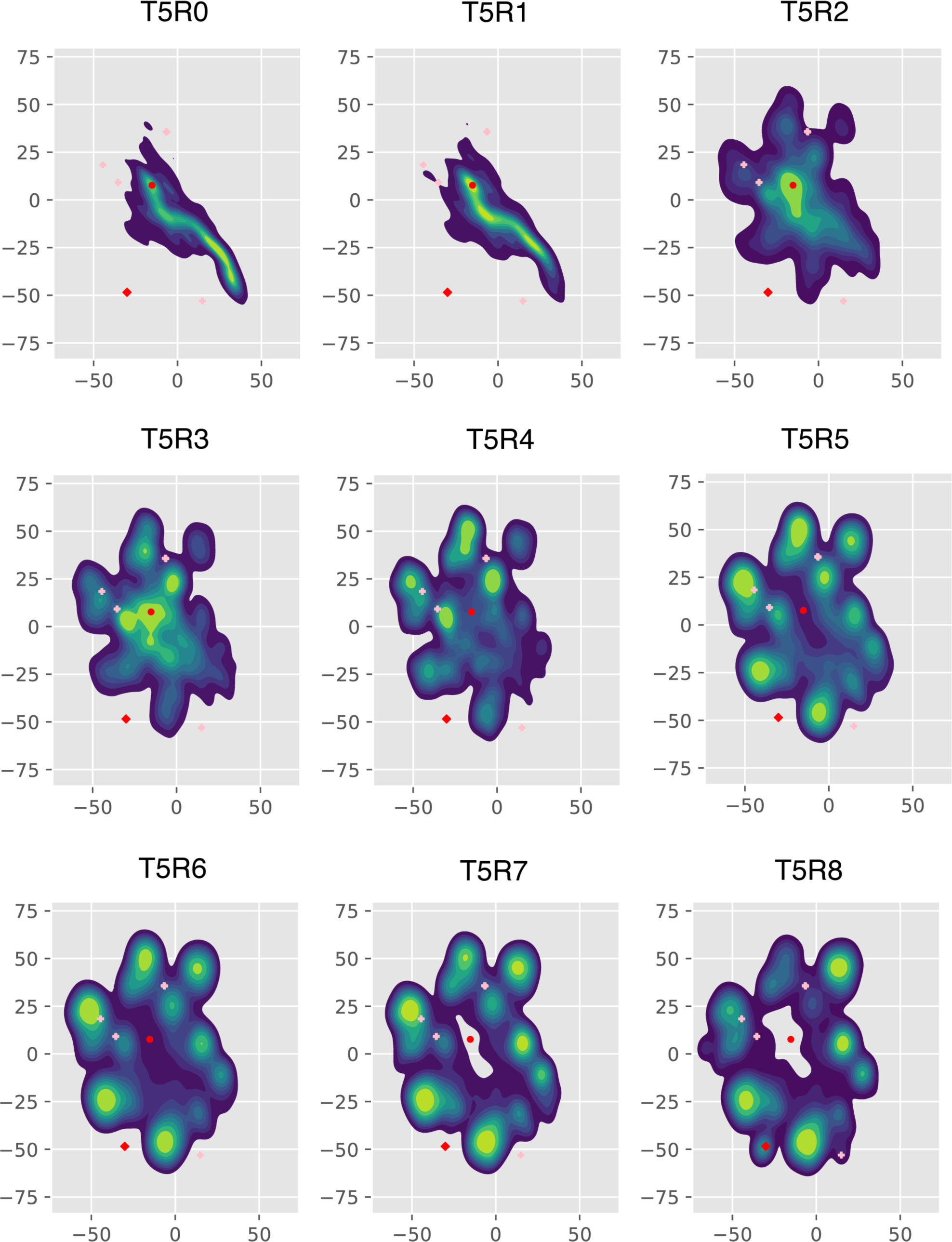
Sequence diversity in T5 selection. Sequence space evolution as shown by t-stochastic neighbour embedding (tSNE^2^) as a two-dimensional projection visualisation across the whole T5 selection trajectory for T5R0-T5R8 (T5 seed sequence (red dot), T5R8/359 aptamer (red diamond), ATP aptamer motif perfect hits (Fig. 2, pink crosses).

**Supplementary Fig. 9.**
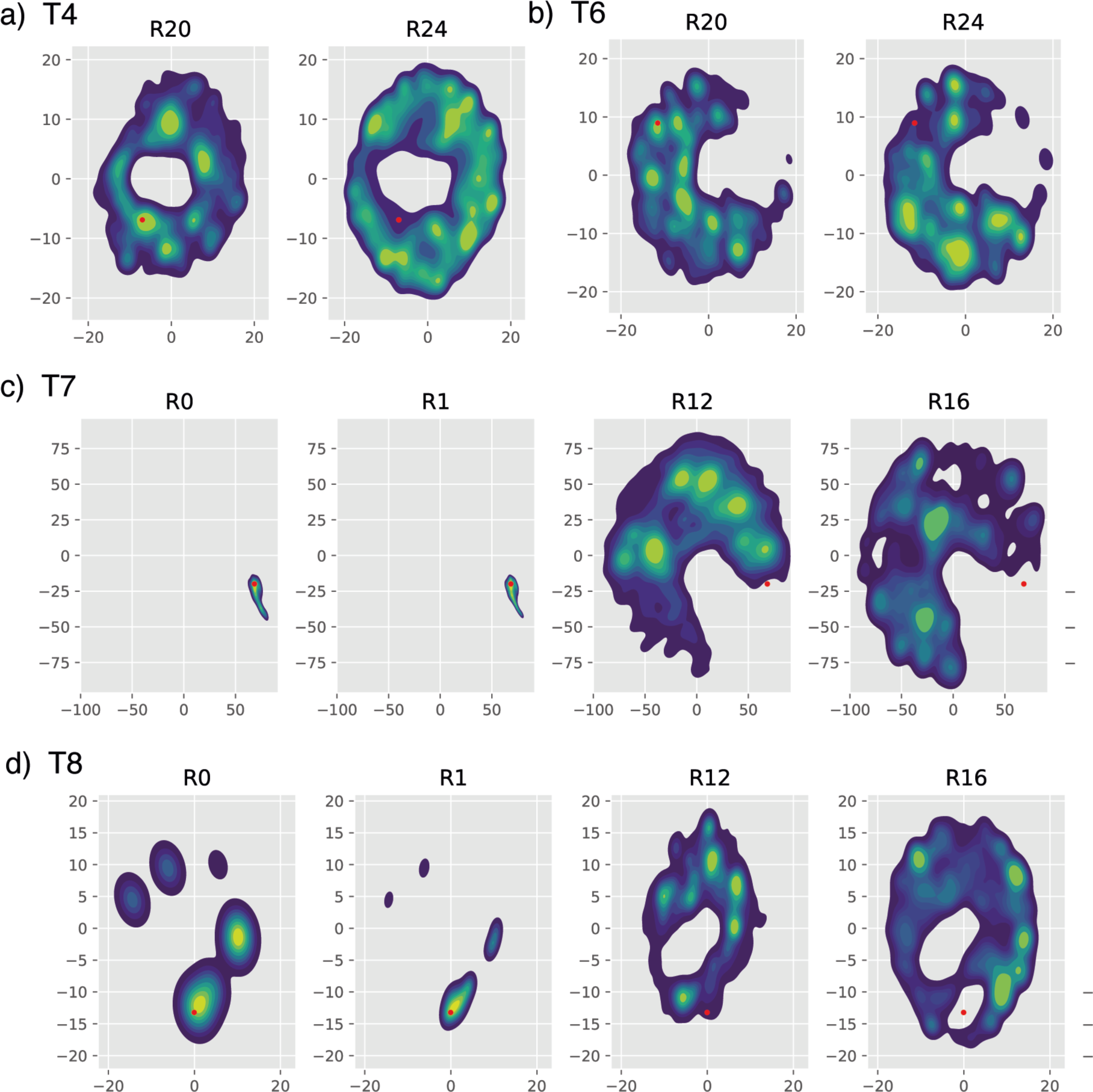
Sequence diversity in T4, T6, T7 and T8 selections. Sequence space evolution as shown by t-stochastic neighbour embedding (tSNE^2^) as a two-dimensional projection visualisation of a), T4, b) T6, c) T7, d) T8 at different points of the selection (R0-R24). T4, T6,T7,T8 seed sequences shown as red dot.

## Supplementary Tables

**Supplementary Table 1:**
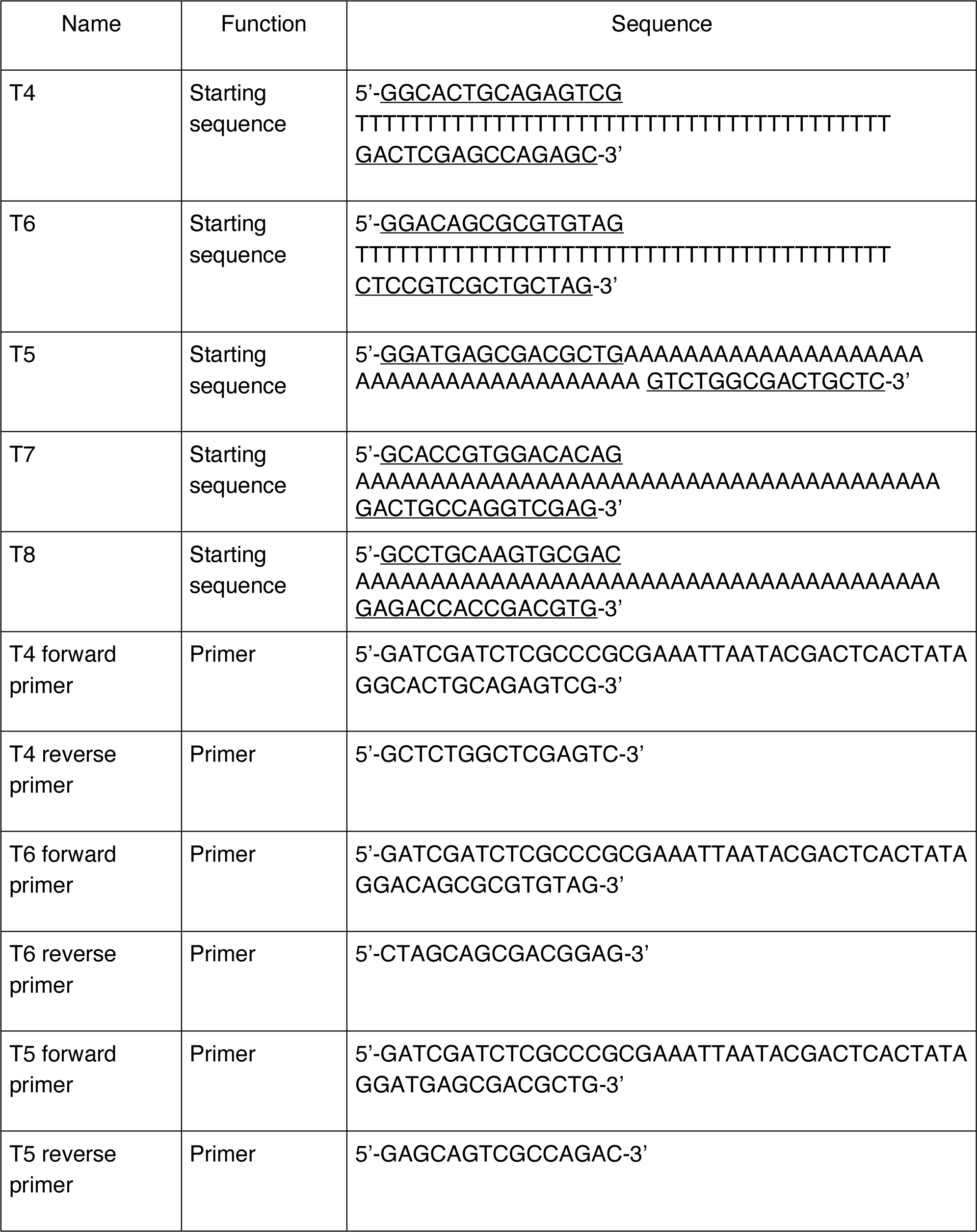
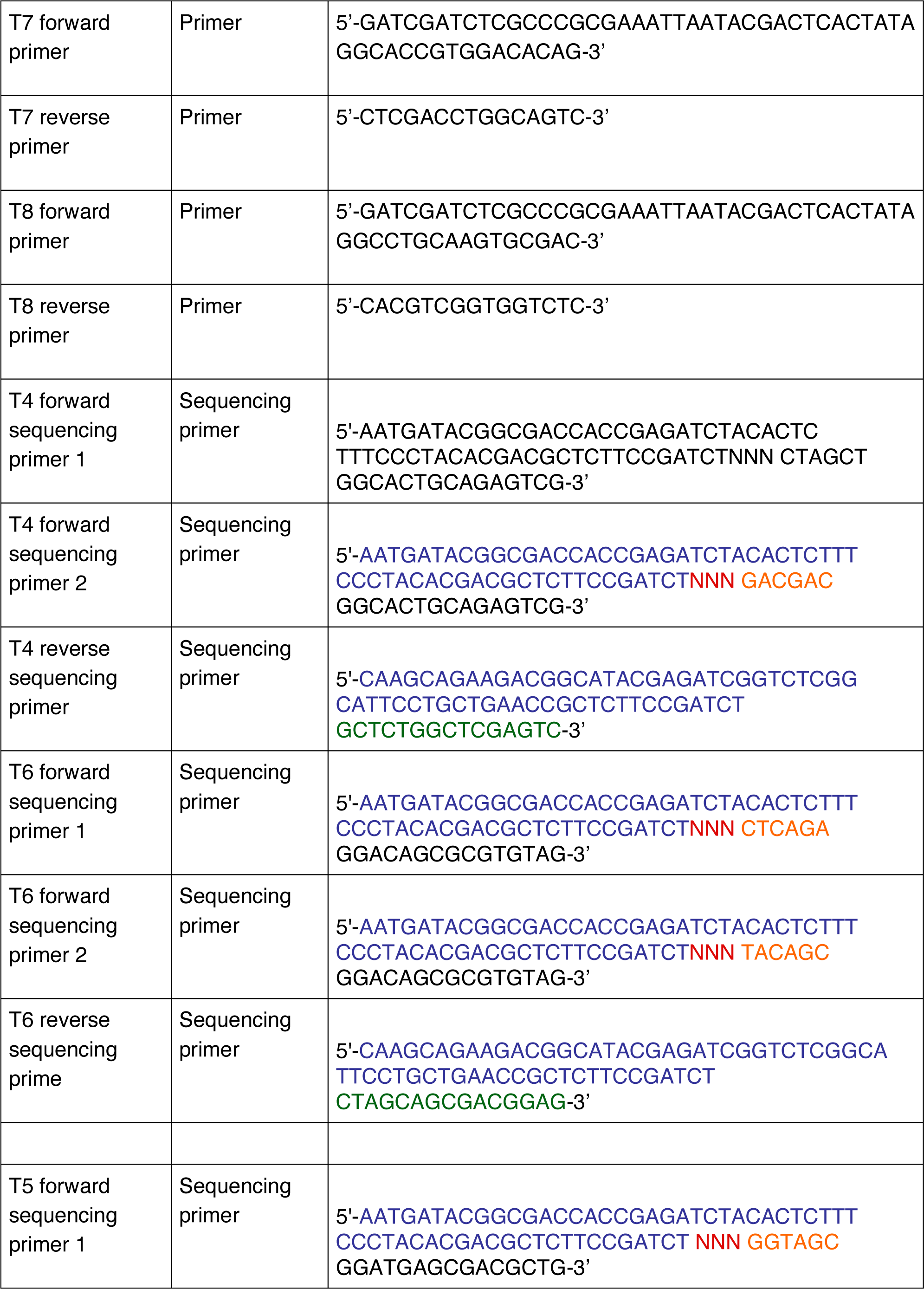
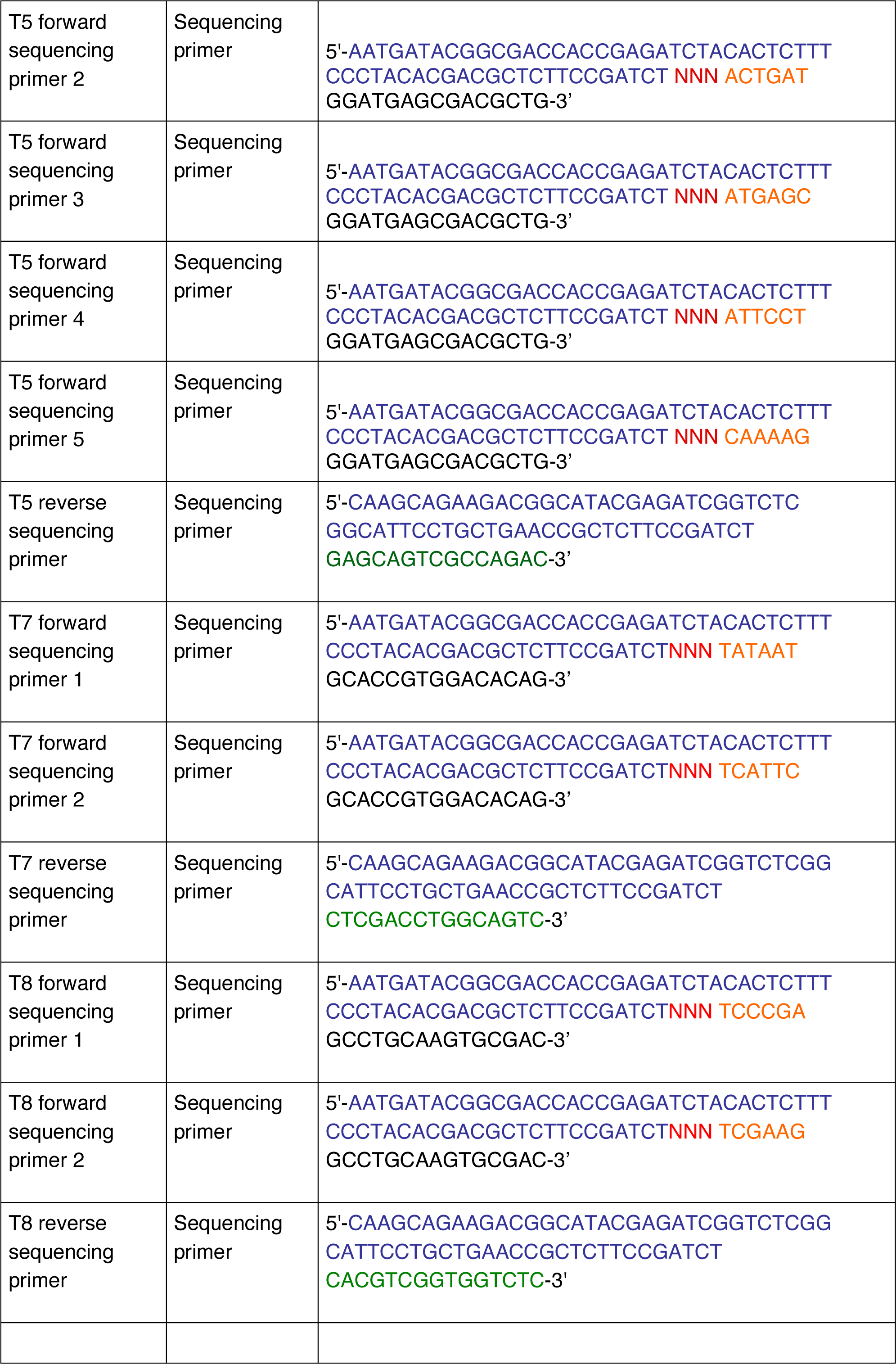
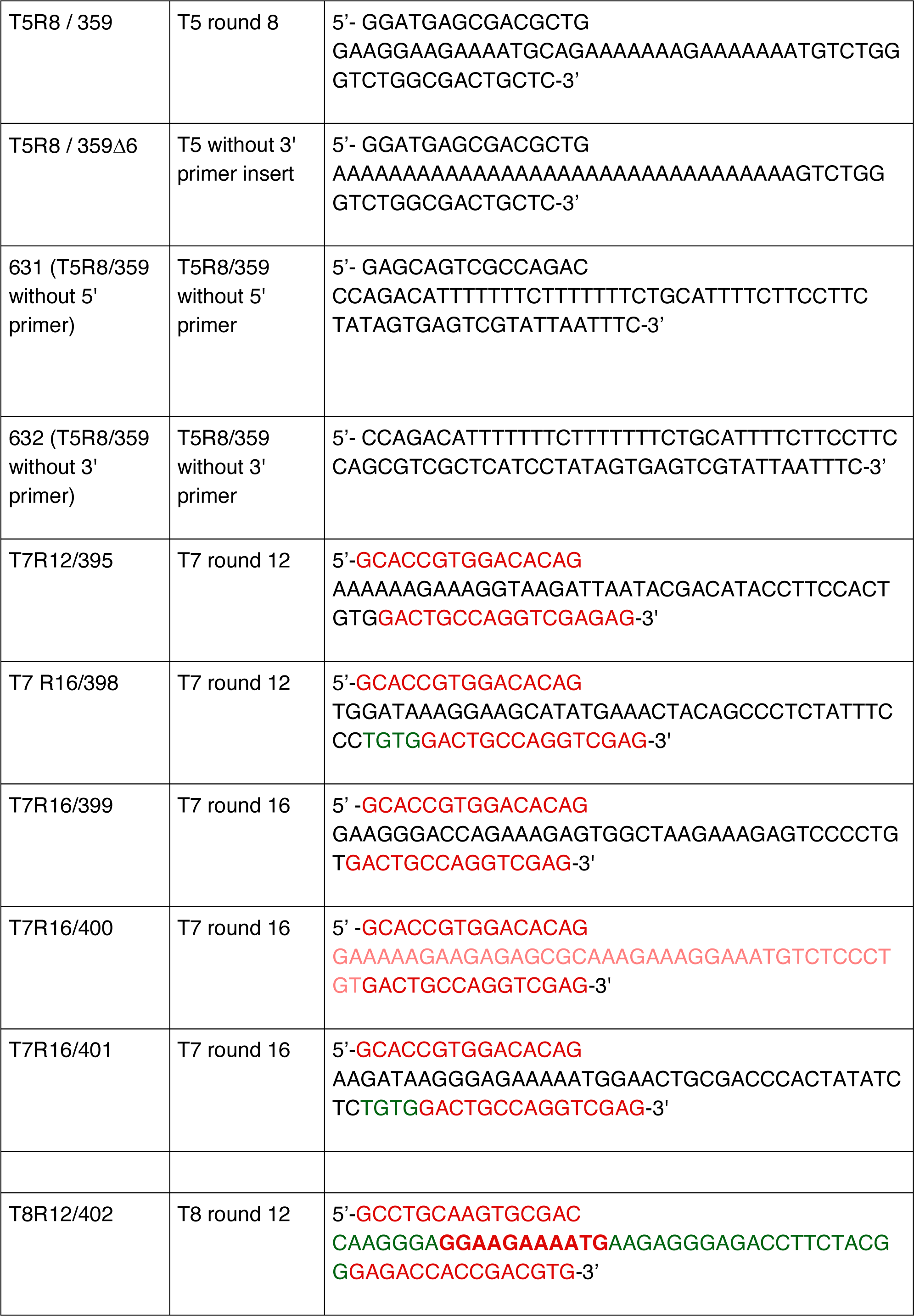
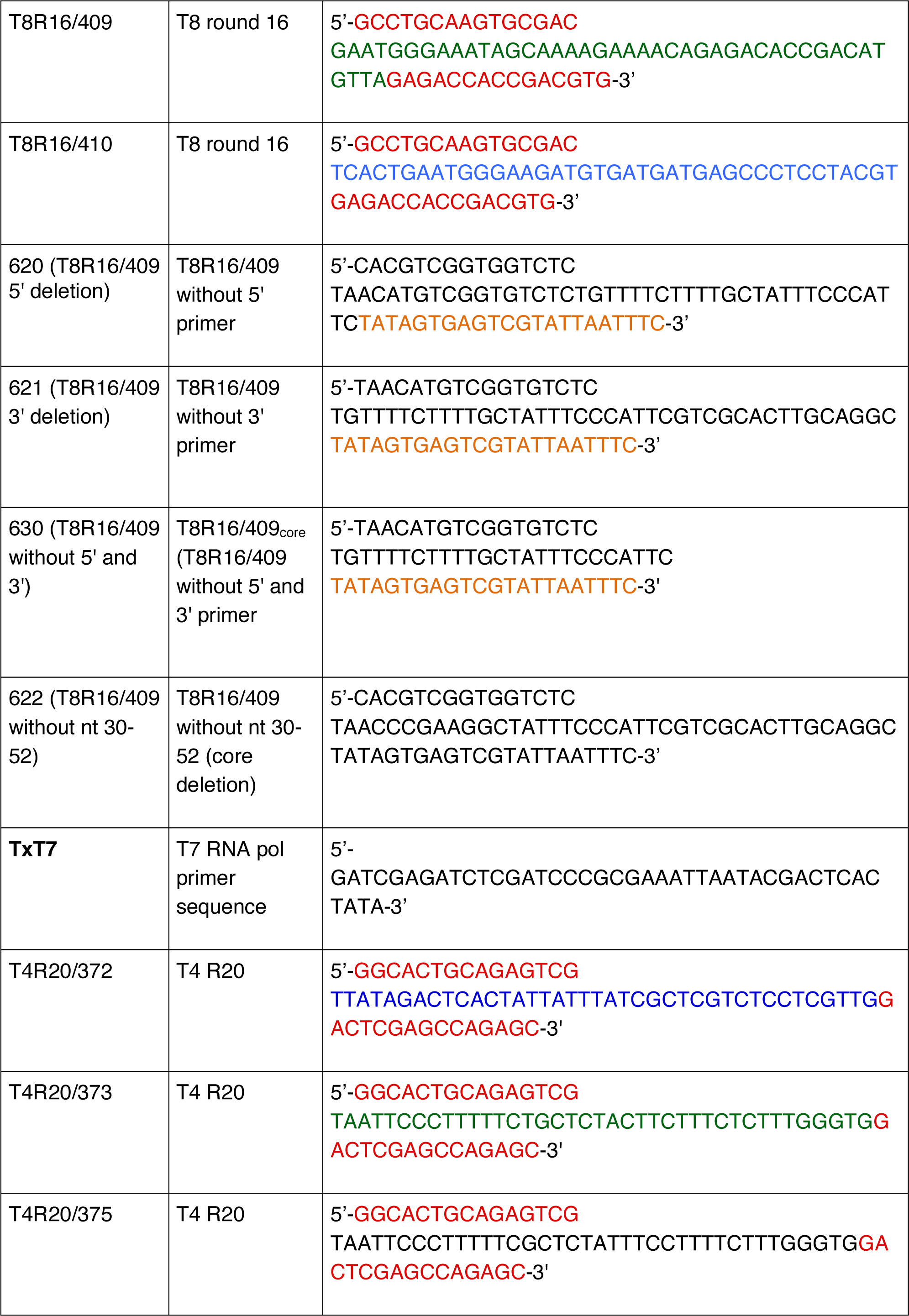
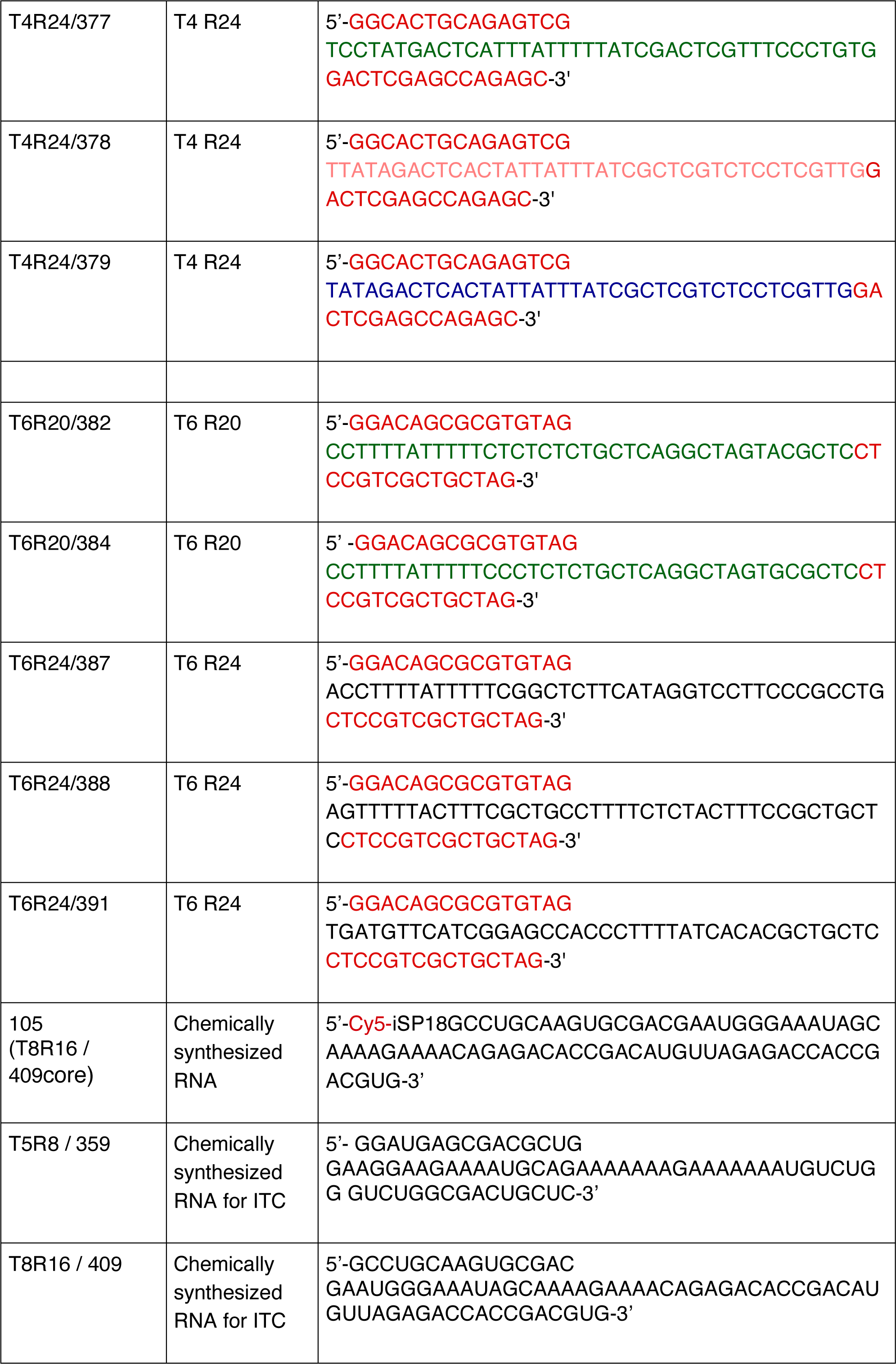
Oligonucleotide sequences. The primer binding sites of the starting sequences are underlined. RNA sequences are shown as DNA templates for PCR. Sequencing barcodes shown in colour.

**Supplementary Table 2.** Column binding and elution assay. Analysis of PAGE gels from column binding assays (ATP agarose) of different RNA sequences (see Supplementary Table 1). Values (in %) represent the amount of RNA for each individual fraction on the gel, data were generated from 2-3 experimental repeats. * Fractions 3-5 are combined.

| | T5 | T5R8 / 359Δ6<br>(T5R8 / 359 without<br>3' primer insert) | T5R8/359 |
| --- | --- | --- | --- |
| unbound | 91.7 | 93.4 | 32.8 |
| wash | 5.0 | 2.1 | 7.5 |
| ATP 1 | 1.0 | 1.6 | 44.6 |
| ATP 2 | 0.9 | 1.1 | 7.6 |
| ATP 3 | 1.3 | 1.8 | 7.5* |

|  | T7 | T7R12/<br>395 | T7R16/<br>401 | T7R16/<br>400 | T7 R16/<br>398 | T7R16/<br>399 |
| --- | --- | --- | --- | --- | --- | --- |
| unbound | 92.0 | 86.5 | 87.5 | 88.3 | 89.0 | 89.4 |
| wash | 4.8 | 4.5 | 2.1 | 2.4 | 2.9 | 5.1 |
| ATP 1 | 1.6 | 4.1 | 4.7 | 3.6 | 2.1 | 2.6 |
| ATP 2 | 1.3 | 4.4 | 3.9 | 3.7 | 3.6 | 1.5 |
| ATP 3 | 0.3 | 0.4 | 1.8 | 2.1 | 2.4 | 1.4 |

|  | T8 | T8R12/<br>402 | T8R16/<br>410 | T8R16/<br>409 | T8R16/<br>409 |
| --- | --- | --- | --- | --- | --- |
| unbound | 94.9 | 84.5 | 91.9 | 35.1 | 32.7 |
| wash | 3.8 | 5.3 | 3.7 | 6.7 | 6.4 |
| ATP 1 | 0.0 | 7.9 | 1.9 | 27.7 | 20.9 |
| ATP 2 | 0.1 | 2.2 | 1.9 | 19.1 | 14.4 |
| ATP 3 | 1.1 | 0.1 | 0.6 | 11.4 | 11.8 |
| ATP 4 |  |  |  |  | 4.6 |
| ATP 5 |  |  |  |  | 5.7 |
| ATP 6 |  |  |  |  | 3.5 |

|  | T4 | T4R20/<br>372 | T4R20/<br>373 | T4R20/<br>375 | T4R24/<br>377 | T4R24/<br>378 | T4R24/<br>379 |
| --- | --- | --- | --- | --- | --- | --- | --- |
| unbound | 92.0 | 87.5 | 90.2 | 95.6 | 84.3 | 93.8 | 89.5 |
| wash | 5.1 | 3.3 | 3.4 | 1.0 | 3.4 | 3.5 | 4.8 |
| ATP 3 | 0.8 | 5.6 | 1.9 | 2.1 | 6.9 | 1.2 | 3.5 |
| ATP 2 | 1.1 | 1.1 | 2.4 | 0.2 | 2.7 | 0.8 | 1.8 |
| ATP 3 | 0.9 | 2.5 | 2.1 | 1.2 | 2.6 | 0.7 | 0.4 |

|  | T6 | T6R20/<br>382 | T6R20/<br>384 | T6R20/<br>387 | T6R24/<br>391 | T6R24/<br>388 |
| --- | --- | --- | --- | --- | --- | --- |
| unbound | 92.3 | 89.2 | 91.3 | 92.9 | 92.3 | 83.3 |
| wash | 3.6 | 4.6 | 3.0 | 3.4 | 5.3 | 2.8 |
| ATP 1 | 1.0 | 3.2 | 2.3 | 2.2 | 1.1 | 5.8 |
| ATP 2 | 0.7 | 1.8 | 2.0 | 0.9 | 0.5 | 6.4 |
| ATP 3 | 2.4 | 1.2 | 1.4 | 0.6 | 0.9 | 1.8 |

**Supplementary Table 3.** Column binding and elution assay of GTP aptamer variants. Analysis of PAGE gels from column binding assays (ATP agarose) of different RNA sequences (see Supplementary Table 1). Values (in %) represent the amount of RNA for each individual fraction on the gel, data were generated from 2-3 experimental repeats. * Fractions 3-5 are combined.

|  | 620<br>(T8R16/409<br>without 5'<br>primer) | 621<br>(T8R16/409<br>without 5'<br>primer) | 630<br>(T8R16/<br>409core) | 622<br>(T8R16/409<br>without core) | 105<br>(T8R16/409<br>core RNA) |
| --- | --- | --- | --- | --- | --- |
| unbound | 47.6 | 47.6 | 59.7 | 90.7 | 35.2 |
| wash | 8.4 | 12.8 | 6.9 | 4.6 | 2.3 |
| ATP 1 | 26.1 | 20.2 | 11.7 | 3.6 | 27.5 |
| ATP 2 | 11.3 | 11.5 | 14.0 | 0.3 | 20.6 |
| ATP 3 | 6.5 | 7.9 | 7.7 | 0.8 | 14.4* |

|  | 105<br>(buffer elution) | 105<br>(ATP elution) | 105<br>(CTP elution) | 105<br>(GTP elution) | 105<br>(UTP elution) |
| --- | --- | --- | --- | --- | --- |
| unbound | 84.2 | 24.1 | 34.5 | 29.9 | 34.7 |
| wash | 5.3 | 7.5 | 6.4 | 6.3 | 9.9 |
| ATP 1 | 3.2 | 29.8 | 18.5 | 46.9 | 9.9 |
| ATP 2 | 2.3 | 20.6 | 16.8 | 9.6 | 21.3 |
| ATP 3 | 1.8 | 9.2 | 10.9 | 3.6 | 14.4 |
| ATP 4 | 1.7 | 5.8 | 7.7 | 2.3 | 3.6 |
| ATP 5 | 1.5 | 3.0 | 5.3 | 1.4 | 6.1 |

## Notes

### Competing Interest Statement

The authors have declared no competing interest.

